# Autophagy is a cell-intrinsic driver of neural stem cell quiescence in hippocampal dentate gyrus development

**DOI:** 10.1101/2023.05.03.539225

**Authors:** Isabel Calatayud-Baselga, Lucía Casares-Crespo, Carmina Franch-Ibáñez, José Guijarro-Nuez, Pascual Sanz, Helena Mira

## Abstract

Neurogenesis in the adult mammalian brain relies on the lifelong persistence of quiescent neural stem cell (NSC) reservoirs. Little is known about the mechanisms that lead to the initial establishment of NSC quiescence during development. Here, we show that protein aggregates and autophagy machinery components accumulate in quiescent NSCs and that pharmacological blockade of autophagy disrupts quiescence. Conversely, increasing autophagy through AMPK/ULK1 activation instructs the acquisition of the quiescent state. Selective ablation of *Atg7*, a critical gene for autophagosome formation, in hippocampal radial-glia like NSCs at early and late postnatal stages compromises the initial acquisition and maintenance of quiescence during the formation of the dentate gyrus SGZ niche. Therefore, we demonstrate that autophagy is cell-intrinsically required to establish radial glia-like NSC quiescence during hippocampal development. Our results uncover a fundamental role of autophagy in the transition of developmental NSCs into their dormant adult form, paving the way for studies directed at further understanding the mechanisms of stem cell niche formation and maintenance in the mammalian brain.

## Introduction

Neurogenesis continues throughout life in the adult mammalian brain thanks to neural stem cells (NSCs) persisting in defined niches beyond development. These stem cells ultimately derive from proliferating developmental precursors that are set aside as reservoirs, either embryonically in the ventricular-subventricular zone (V-SVZ) niche^1,2^ or postnatally in the hippocampal dentate gyrus subgranular zone (SGZ) niche^3,4^. Acquisition of quiescence (a reversible exit from the cell cycle) while maintaining stemness is considered a critical turning point that defines the conversion of developmental stem cells to their adult counterparts^5,6^. In adulthood, quiescence is maintained by dominant extrinsic signals produced by various mature niche cell types^7,8^. NSC quiescence is initially established before a fully functional adult niche structure is assembled, so it is currently unknown how NSCs first come to rest^6^.

Using the hippocampal dentate gyrus as a model system, we hypothesised that cell-intrinsic mechanisms, including those related to proteostasis, could contribute to quiescence entry during development. Proteostasis is key for the fitness of long-lasting cells like stem cells^9^. It controls protein turnover through intertwined pathways regulating protein synthesis, folding and degradation through the ubiquitin–proteasome system (UPS) or the autophagy–lysosomal pathway (ALP). Previous studies focused on adult NSCs reported lower protein synthesis, protein aggregate accumulation and higher lysosomal activity in adult quiescent NSCs acutely isolated from mouse V-SVZs^10^; FOXO3 regulation of a network of autophagy genes in cultured adult V-SVZ NSCs^11^ ; a role for aggresomes in the clearance of protein aggregates during adult NSC activation^12^; and increased adult SGZ NSC proliferation in conditional knockouts of the lysosomal master regulator *Tfeb*^13^. Together, the aforementioned results in adult NSCs suggest that ALP could be playing an additional, unexplored, cell-autonomous function in driving the initial establishment of quiescent NSC reservoirs during niche development. To tackle this question, we focused specifically on the possible role of autophagy in the acquisition of quiescence by radial glia–like NSCs located in the hippocampal SGZ niche, a poorly understood developmental transition.

## Results

Embryonic dentate progenitors migrate from the neuroepithelium to the primitive dentate gyrus, forming the definitive adult radial glia–like NSC (RGL) reservoir during early postnatal development^3,4,14^. After a proliferation peak at postnatal day 3 (P3), RGLs occupy their final position in the SGZ at around P14 and continue to enter quiescence until P21^3,4^ (Extended Data Fig. 1 a-d). We first set out to measure overall proteostatic events in P3 and P21 hippocampal NSCs at the single-cell level. NSCs were acutely isolated by FACS on the basis of the combined expression of the cell surface markers GLAST and Prominin-1^15,16^ (Fig. 1a, Extended Data Fig. 2a). We compared GLAST^+^PROM-1^+^ cells at both ages. P21 NSCs substantially accumulated protein aggregates (visualized by the trapping of the fluorescent molecular rotor ProteoStat; Fig. 1a-b), a finding corroborated in hippocampal GLAST^+^ cells isolated by MACS (Extended Data Fig. 2b-d). At P21, proliferation of GLAST^+^ cells (percentage of Ki-67^+^ cells) markedly decreased (Fig. 1c) and global protein synthesis (OP-Puro signal) was reduced (Fig. 1d), but protein aggregates (ProteoStat signal, Fig. 1e) and lysosomes (LAMP2 signal, Fig. 1f) accumulated, pointing to a shift in hippocampal RGL proteostasis during the P3 (proliferation) to P21 (quiescence) developmental transition. Likewise, quiescent P21 and P14 RGLs identified *in vivo* as SOX2/GFAP co-expressing cells with radial morphology that were negative for the cell cycle marker Ki-67, showed increased LAMP2 levels compared to actively dividing (Ki-67^+^) RGLs (Fig. 1g-h).

**Figure 1.**
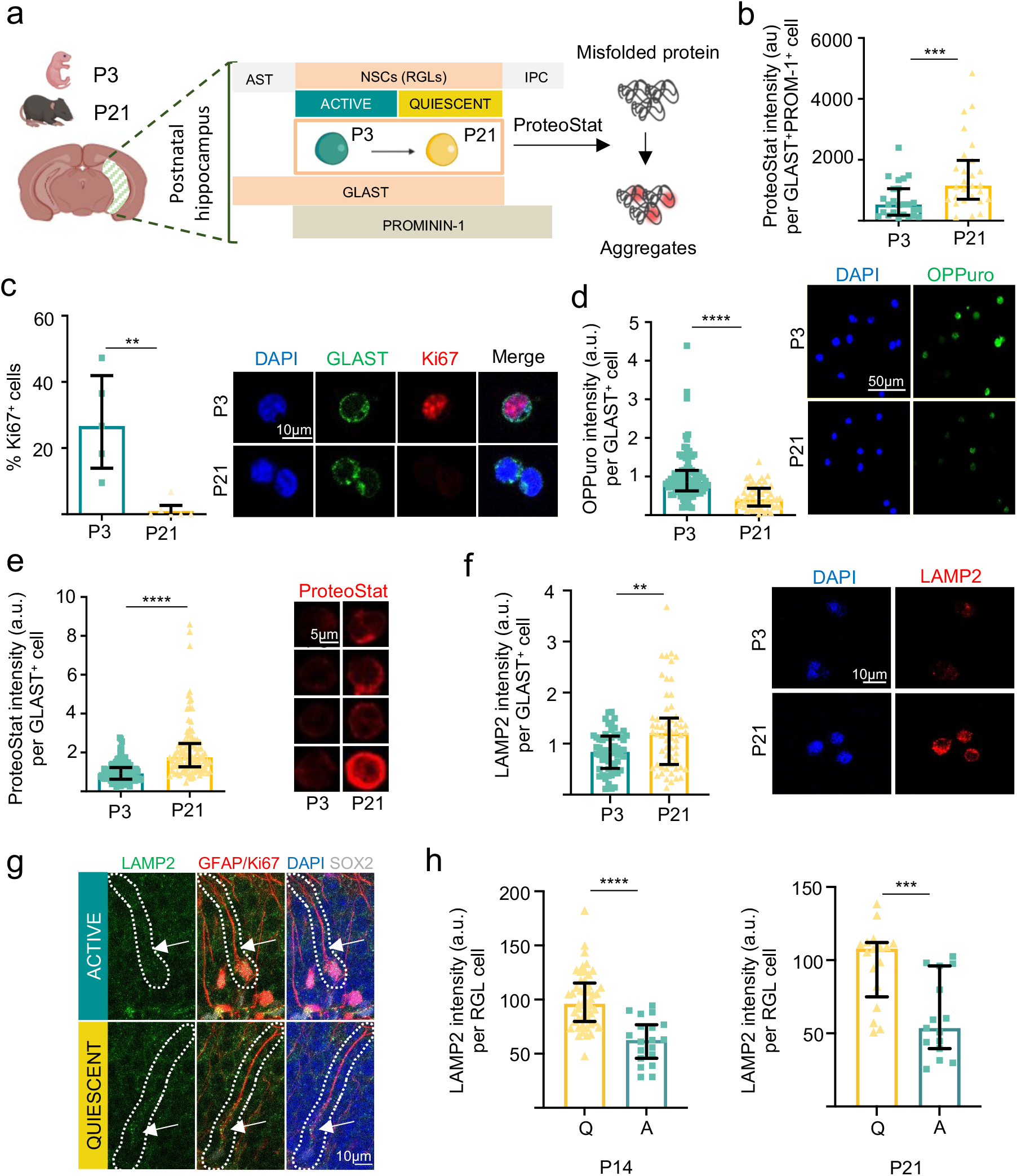
Protein aggregates and lysosomes accumulate in quiescent P21 hippocampal NSCs compared to proliferating P3 NSCs. **a**. Experimental design. ProteoStat labelling of protein aggregates was performed on GLAST^+^Prominin-1^+^ NSCs isolated by FACS from the hippocampus of mice on postnatal day 3 (P3) and P21. AST, astrocyte. IPC, intermediate progenitor cell. **b**. Mean fluorescence of ProteoStat per GLAST^+^Prominin-1^+^ per cell, measured by confocal microscopy. Each point represents an analyzed cell. **c**. (Left) Percentage of proliferative cells (Ki67^+^) cells isolated from P3 and P21 mice. (Right) Representative immunocytochemistry images of GLAST/Ki67 in GLAST^+^ cells isolated of postnatal 3 and 21-day-old mice. **d**. (Left) Quantification of OPPuro fluorescence in GLAST^+^ cells isolated from P3 and P21 old mice. (Right) Representative immunofluorescence confocal images of GLAST^+^ cells isolated from P3 and P21 mice treated with OPPuro, which is incorporated in newly synthesized proteins. **e**. (Left) Quantification of protein aggregates measured with ProteoStat in GLAST^+^ cells. (Right) Representative immunofluorescence confocal images of GLAST^+^ cells isolated from P3 and P21 mice stained with ProteoStat (red). **f**. (Left) Quantification of LAMP2 in GLAST^+^ cells. (Right) Representative immunofluorescence confocal images of GLAST^+^ cells isolated from P3 and P21 mice stained with LAMP2. **g**. (Left) Representative immunohistochemistry confocal images of LAMP2 (green) in active and quiescent RGLs labelled for GFAP (red), SOX2 (white) and Ki67 (red). **h**. Quantification of LAMP2 intensity in active (Ki67^+^) or quiescent (Ki67^-^) RGLs (GFAP^+^/SOX2^+^) in the dentate gyrus of postnatal 14 (P14, left) and 21 (P21, right) days old wild type mice. Each point represents an analyzed RGL cell. Data in this figure are presented as median ± interquartile range from n≥3 independent isolations. At least 15 cells were analyzed per experiment and condition. Statistics: Mann Whitney test. *p < 0.05; **p<0.01; ***p<0.001; ****p<0.0001. Representation shown in a. was created with BioRender.com.

**Figure 2.**
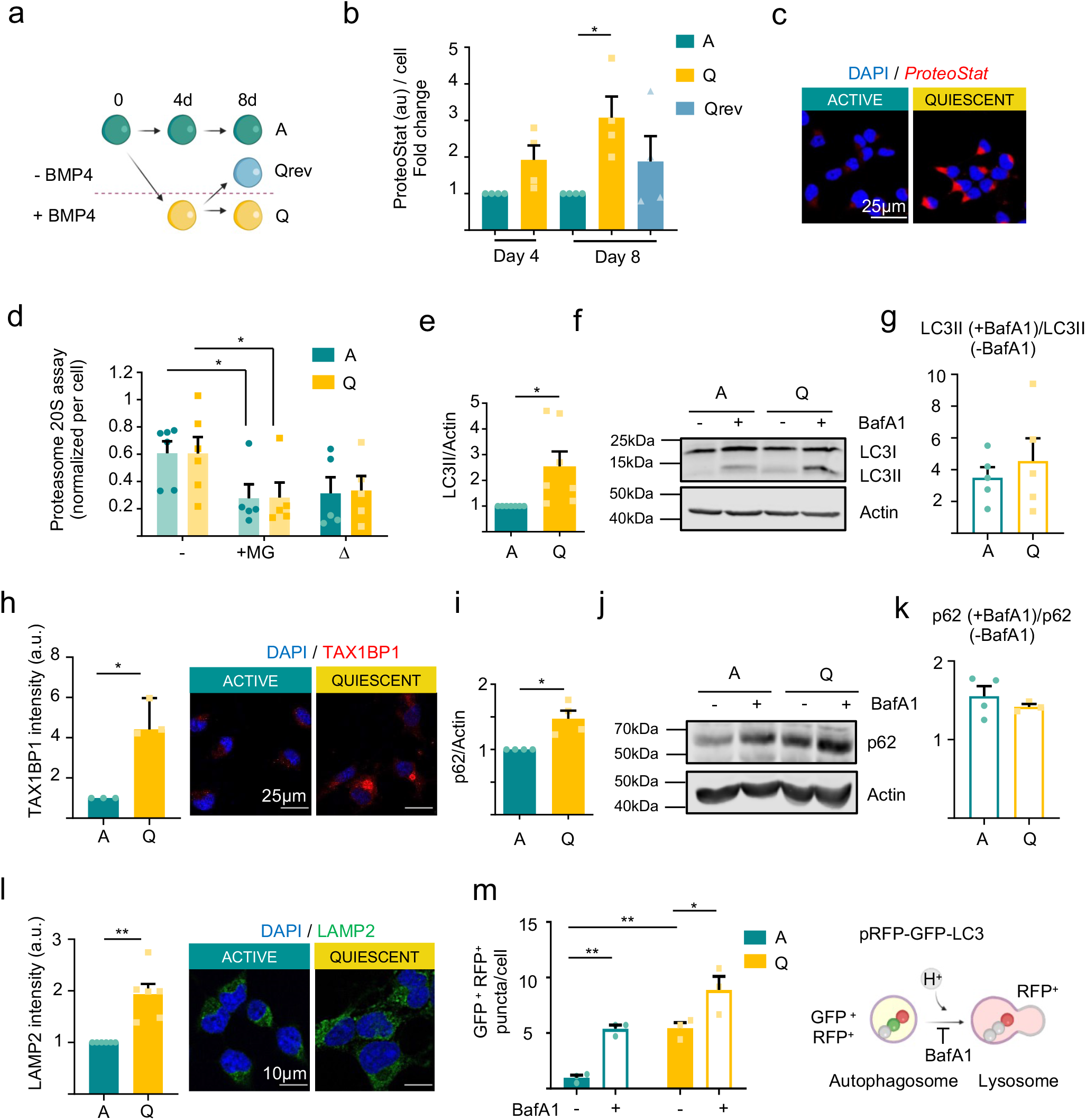
Protein aggregates accumulate in quiescent NSPC cultures despite the increase in autophagy machinery content. **a**. Experimental design. Hippocampal NSPC cultures are grown in the presence of fibroblast growth factor 2 (FGF2) and quiescence is induced by the addition of bone morphogenetic protein 4 (BMP4). Removing BMP4 reverses quiescence. A, active NSPCs; Q, quiescent NSPCs; Qrev, quiescence-reverted NSPCs. **b**. Quantification of protein aggregates measured by ProteoStat labelling at 4 and 8 days of quiescence induction and 4 days of quiescence reversion. At least 15 cells were analyzed per experiment and condition. Each point represents a single cell. Statistics: Day 4, one-sample t-test. **c**. Representative confocal immunofluorescence images of Q-NSPC and A-NSPC cultures stained with ProteoStat (red). **d**. Quantification of proteasome 20S chymotrypsin activity (Δ) in A- and Q-NSPCs upon its inhibition using 10 μM MG132. Statistics: two-way ANOVA. **e**. Quantification of LC3-II levels in A- and Q-NSPCs. Statistics: one-sample t-test. **f**. Immunoblots of LC3 in A- and Q-NSPCs after 100 nM BafA1 treatment (6 h). β-actin was used as a loading control. **g**. The autophagy flux expressed as the ratio of LC3-II(+BafA1)/LC3-II(-BafA1) was equivalent for A-NSPCs and Q-NSPCs. **h**. (Left) Quantification of TAX1BP1 protein level measured as fluorescence intensity. Each point represents individual cultures. Statistics: one sample t-test. (Right) Representative immunofluorescence confocal images of TAX1BP1 in A- and Q-NSPCs. **i**. Quantification of p62 levels in A- and Q-NSPCs. Statistics: one-sample t-test. **j**. Immunoblots of p62 in A- and Q-NSPCs after 100 nM BafA1 treatment (6 h). β-actin was used as a loading control. **k**. Quantification of p62 accumulation after BafA1 treatment in A- and Q-NSPCs. Statistics: one-sample t-test and unpaired t-test, respectively. **l**. (Left) Quantification of LAMP2 protein level measured as fluorescence intensity. Each point represents individual cultures. Statistics: one sample t-test. (Right) Representative immunofluorescence confocal images of LAMP2 in A- and Q-NSPCs. **m**. (Right) In electroporated cells expressing the pRFP-GFP-LC3 sensor, autophagosomes display GFP and RFP fluorescence (yellow puncta), whereas autolysosomes display RFP fluorescence only, because GFP is denatured at the acidic pH of the lysosome. (Left) Quantification of the number of autophagosome puncta (GFP^+^RFP^+^) in A- and Q-NSPCs after 100 nM BafA1 treatment (6 h). Each point represents a single cell. Statistics: two-way ANOVA. Data are presented as mean ± SEM from n≥3 independent cultures. *p < 0.05; **p<0.01; ***p<0.001. Representations shown in a., and n. were created with BioRender.com.

To explore the functional significance of these observations, we moved to an *in vitro* system that allows the controlled induction of quiescence in hippocampal neural stem and progenitor cell (NSPC) cultures, employing the quiescence-promoting signal BMP4^17,18^ (Fig. 2a). In line with the FACS/MACS data, protein aggregates accumulated in quiescent (Q) NSPCs compared to active (A) NSPCs (Fig. 2b-c). When quiescence was reverted by BMP4 removal (Qrev), protein aggregate levels partially returned to those found in the A state (Fig. 2b). We then analysed the activity of the two main branches of protein catabolism (UPS and ALP) to investigate if aggregate accumulation was due to a protein degradation failure. We found no differences in proteasome activity between active (A) and quiescent (Q) NSPCs (∆, Fig. 2d). However, Q-NSPCs accumulated LC3-II and SQSTM/p62 (autophagy markers; Fig. 2e-g and Fig. 2i-k, respectively, see also Extended Data Fig. 3a-b and Extended Data Fig. 4a-c) along with Tax1bp1 (aggrephagy receptor; Fig. 2h) and LAMP2 (lysosomal marker; Fig. 2l). Thus, we next evaluated autophagy flux, tracking the LC3-II increase upon disruption of autophagosome–lysosome fusion and lysosome acidification with Bafilomycin A1 (BafA1)^19^. The LC3-II/ β-actin increase in BafA1-treated cells vs. BafA1-untreated cells was similar in both cellular states (Fig. 2e-g) despite a higher initial LC3-II level in Q-NSPCs (Fig. 2e; see Extended Data Fig. 3 for LC3-II levels normalized to total protein). As shown in Fig. 2g, the autophagy flux expressed as the ratio of LC3-II(+BafA1)/LC3-II(-BafA1) was equivalent for A-NSPCs and Q-NSPCs. Similar results were obtained when SQSTM/p62 dynamics were evaluated as a proxy for the autophagy flux (Fig. 2k; see Extended Data Fig. 4 for p62 levels normalized to total protein and for p62 levels quantified per cell by immunofluorescence). To further assess the transition from autophagosomes to autolysosomes, we electroporated the NSPCs with a pRFP-GFP-LC3 tandem sensor^20^, which expresses the red fluorescent protein (RFP) fused to LC3 and a pH-sensitive green fluorescent protein (GFP). This sensor allows autophagosomes to be identified as yellow (green and red) cytoplasmic puncta, whereas autolysosomes are visualized as red puncta, having lost the GFP signal (Fig. 2m). In line with our previous results, Q-NSPCs had more GFP^+^RFP^+^ autophagosomes than A-NSPCs after BafA1 treatment, but a similar autophagic flux (Fig. 2m, Extended Data Fig. 5). These findings suggest that the increase in ALP markers in quiescent cells is likely due to an increase in autophagy machinery content (rather than a deficient autophagic flux), possibly occurring as a compensatory response to the accumulation of protein aggregates.

**Figure 3.**
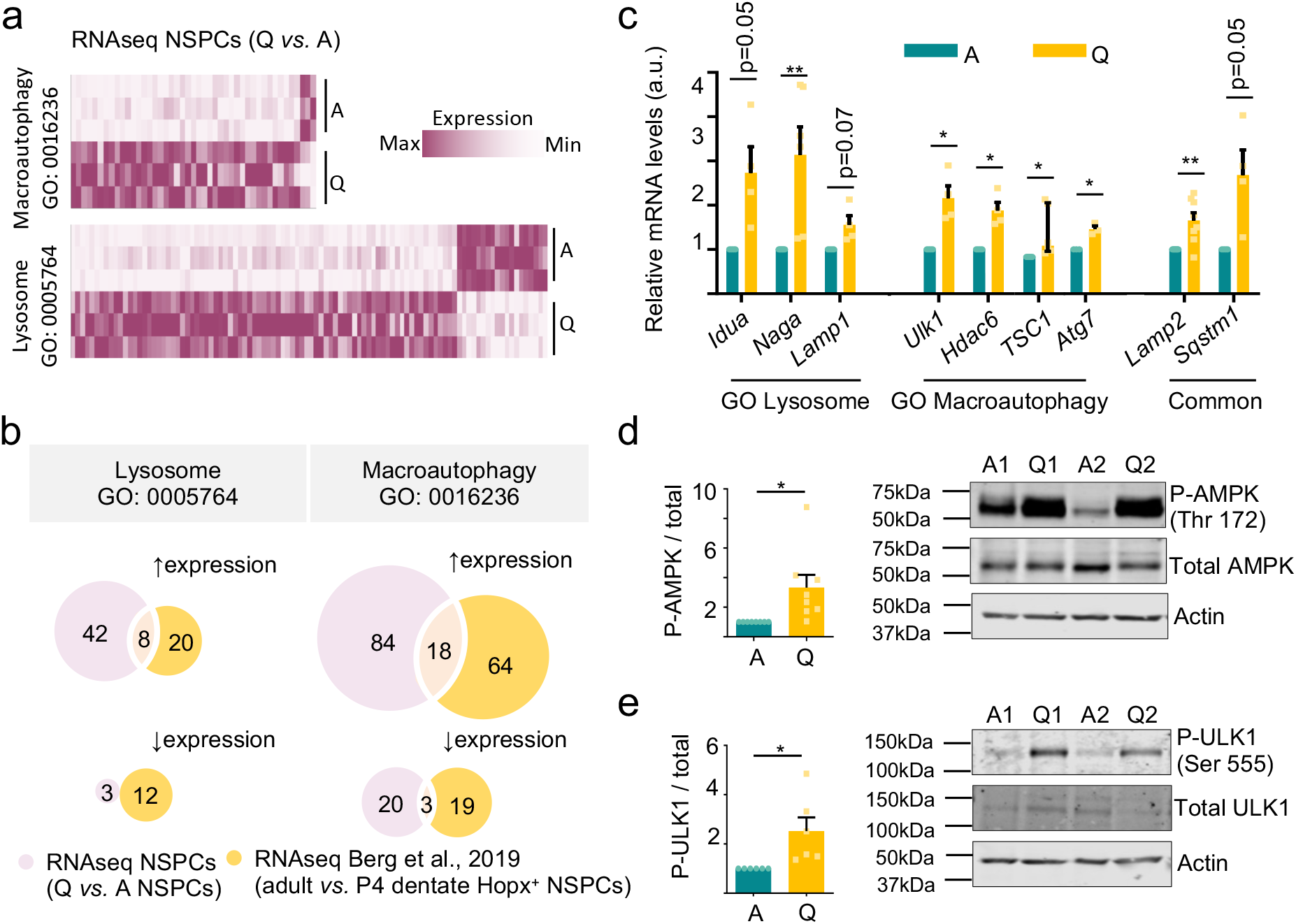
Autophagy is induced in quiescent NSPCs. **a**. Heatmap showing the enrichment in Macroautophagy-GO:0016236 (top) and Lysosome-GO:0005754 (bottom) gene expression in Q-NSPCs versus A-NPSCs. Maximum expression: purple; minimum expression: white. Data were obtained by bulk RNAseq. **b**. Venn diagrams identify common genes associated to Lysosome-GO (Left) and Macroautophagy-GO (Right) in the RNAseq data from this study and RNAseq data from Berg et al., 2019, corresponding to NSCs isolated from 4-day-old (P4) and adult *Hopx-CreER*^*T2*^*::EYFP* mice. NSCs from adult mice are mostly quiescent; P4 NSCs are mostly active. **c**. Relative expression of autophagy–lysosome genes in A- and Q-NSPCs by RT-qPCR analysis. **d**. Left: quantification of phosphorylated P-AMPK (Thr172) levels in A- and Q-NSPCs. Right: representative immunoblots of total AMPK and P-AMPK (Thr172) in A- and Q-NSPCs. β-actin was used as a loading control. **e**. Left: quantification of phosphorylated (P) ULK1 (Ser555) levels in A- and Q-NSPCs. Right: immunoblots of total ULK1 and P-ULK1 (Ser555) in A- and Q-NSPCs. β-actin was used as a loading control. Data are presented as mean ± SEM from n≥3 experiments except for TSC1 in Figure 3c, represented as median ± interquartile range. Statistics: one-sample t-test. *p<0.05; **p<0.01.

**Figure 4.**
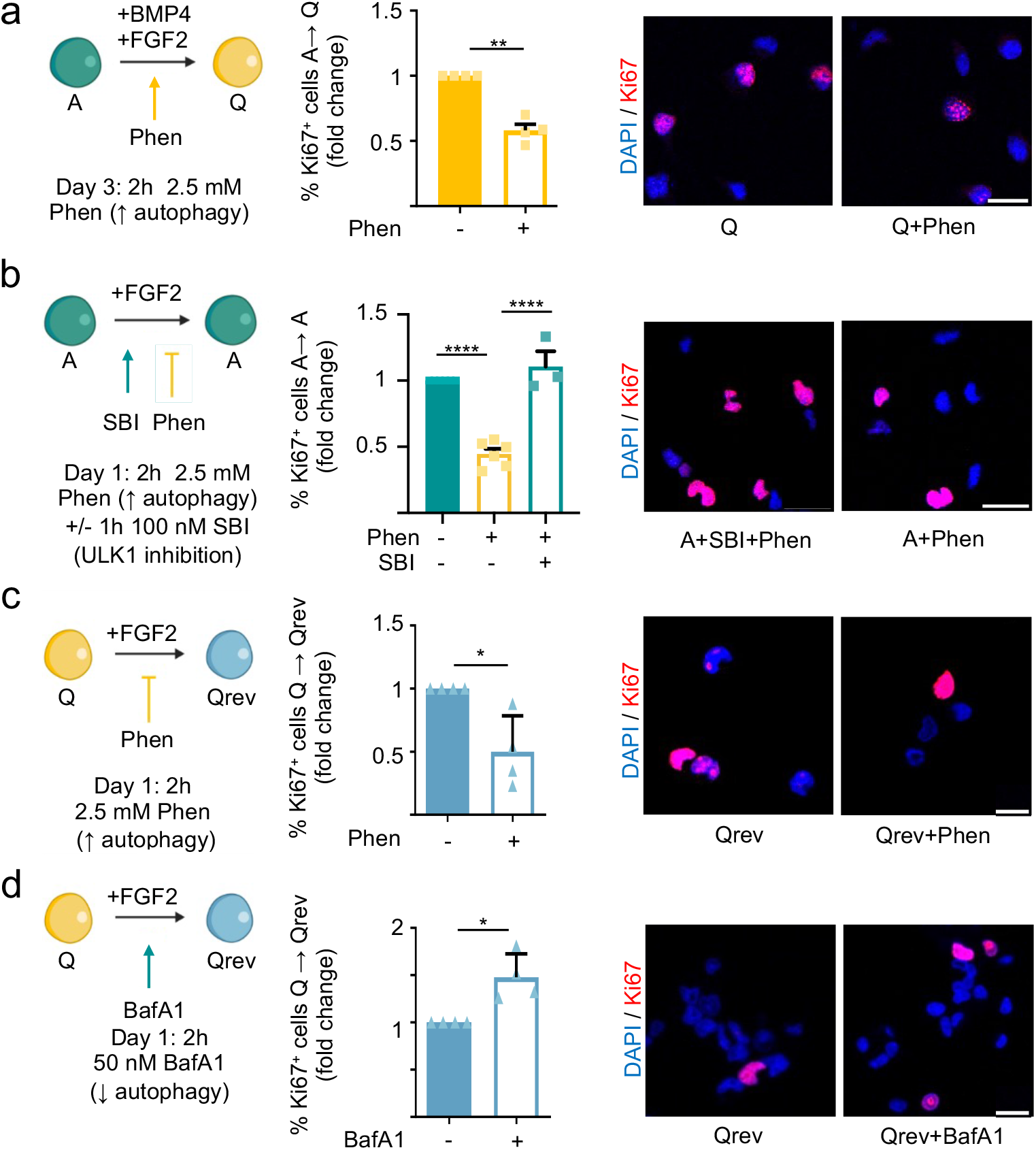
Pharmacological modulation of autophagy regulates the switch between NSPC activation and quiescence. **a**. (Left) Functional assay scheme. Phenformin (Phen) activates the autophagy–lysosome pathway. A-NSPCs were treated for 2 h with Phen on Day 3 of quiescence induction. (Middle) Fold-change in the percentage of Ki-67^+^ cells 24 h after the 2 h Phen 2.5 mM treatment of A-NSPCs entering quiescence. (Right) Representative confocal immunofluorescence images for Ki-67 (red) 24 h after Phen treatment. **b**. (Left) Functional assay scheme. A-NSPCs were treated for 1 h with SBI or DMSO and afterwards 2 h with Phen on Day 1. (Middle) Fold-change in the percentage of Ki-67^+^ cells 24 h after the +/-SBI (100 nM) and +/-Phen treatment (2.5 mM) of A-NSPCs entering quiescence. (Right) Representative confocal immunofluorescence images for Ki-67 (red) 24 h after SBI and Phen treatment. **c**. (Left) Functional assay scheme. (Middle) Fold-change in the percentage of Ki-67^+^ cells 48h after the 2 h Phen (2.5 mM) treatment of Q-NSPCs that were being reverted to activation. (Right) Representative confocal immunofluorescence images for Ki-67 (red) 48 h after Phen treatment. **d**. Bafilomycin A1 (BafA1) inhibits the autophagy–lysosome pathway. (Left) Functional assay scheme. (Middle) Fold-change in the percentage of Ki-67^+^ cells 16 h after the 2 h BafA1 50 nM treatment of Q-NSPCs that were being reverted to activation. (Right) Representative confocal immunofluorescence images for Ki-67 (red) 16 h after BafA1 treatment. Scale bars, 25 μm. Data are presented as mean ± SEM from n≥3 experiments. Statistics: one-sample t-test except Figure 4b: one-way ANOVA was performed. *p<0.05; **p<0.01; ****p<0.0001. Representation shown in a., b., c. and d. were created with BioRender.com.

**Figure 5.**
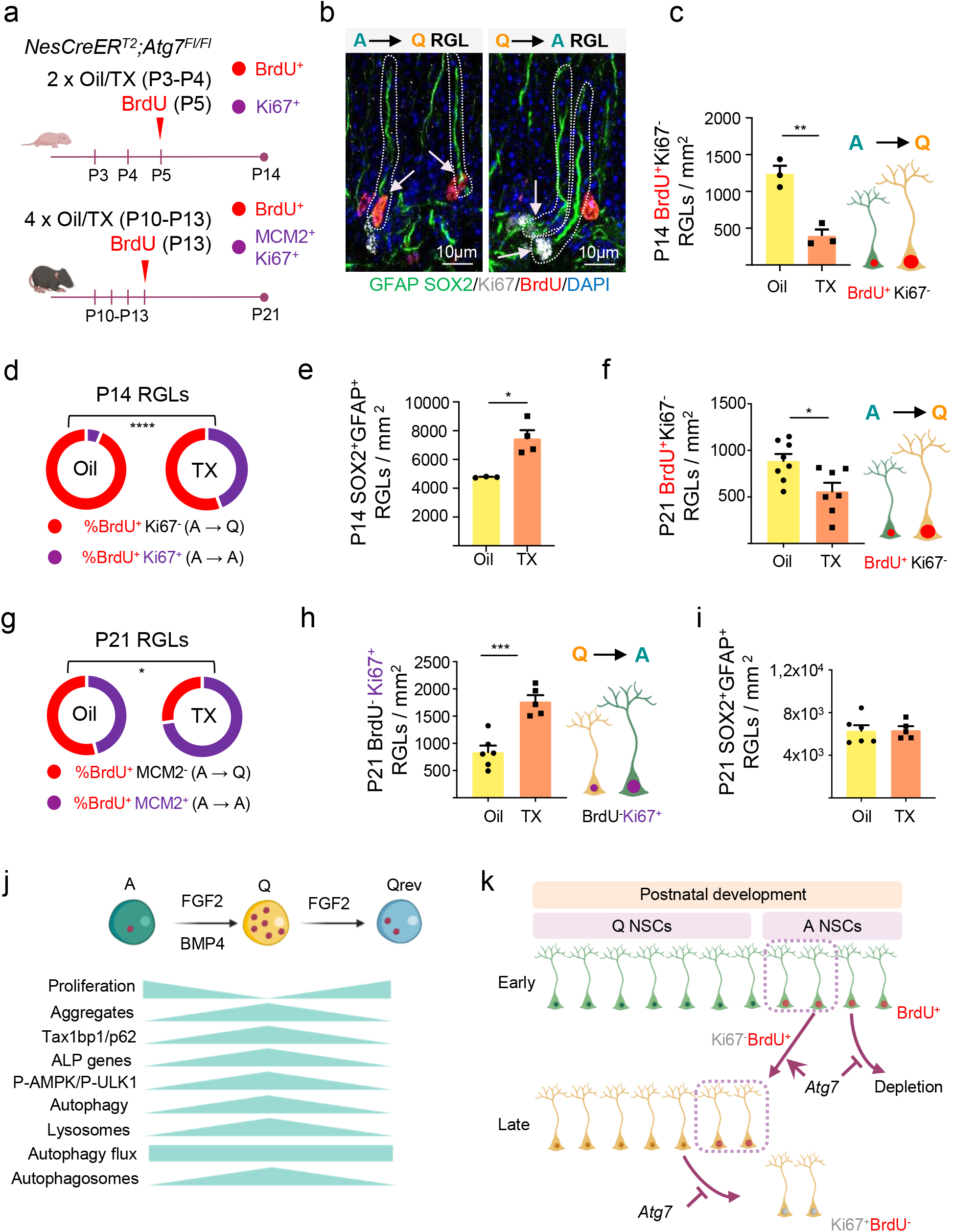
The autophagy gene *Atg7* is cell-autonomously required for the establishment of hippocampal radial glia–like neural stem cell (RGL) quiescence during postnatal development. **a**. Schematic diagram of the experimental design. (Upper pannel) *NesCreER*^*T2*^*;Atg7*^*Fl/Fl*^ mice were injected with tamoxifen (TX) at P3–P4 to induce *Agt7* deletion in NSCs. Oil was used as control. Mice also received BrdU injection to mark dividing cells at P5. Animals were sacrificed at P14 and proliferating cells were identified by Ki-67 staining. (Lowe pannel) *NesCreER*^*T2*^*;Atg7*^*Fl/Fl*^ mice were injected with TX at P10–P13 to induce *Agt7* deletion in NSCs. Oil was used as control. The mice also received BrdU injection to mark dividing cells at P13. Animals were sacrificed at P21 and proliferating cells were identified by Ki-67 or MCM2 staining. **b**. Double-labelling at the time of sacrifice enables the identification of active (A) RGLs that have entered quiescence (Q) (BrdU^+^Ki-67^−^ A→Q RGLs, arrowheads in right panel) and quiescent RGLs that have entered the cell cycle (BrdU^−^Ki-67^+^ Q→A RGLs, arrowheads in left panel). RGLs were identified as SOX2^+^ cells with their soma in the SGZ and a radial GFAP+ process crossing the granule cell layer. Representative confocal images of RGLs labelled for GFAP (green), SOX2 (green), Ki-67 (white) and BrdU (red) at P21 are shown. **c**. Quantification of active RGLs entering quiescence (BrdU^+^Ki-67^−^ RGLs/mm^2^) in the SGZ of *Atg7* conditional knockout (cKO; TX) and control (Oil) P14 mice. **d**. Percentage of P5 active RGLs (BrdU^+^) entering quiescence (BrdU^+^Ki-67^−^, red) or remaining active (BrdU^+^Ki-67^+^, purple) in the SGZ of *Atg7* cKO (TX) and control (Oil) P14 mice. **e**. Quantification of total RGLs (SOX2^+^GFAP^+^ RGLs/mm^2^) at P14. **f**. Quantification of active RGLs entering quiescence (BrdU^+^Ki-67^−^ RGLs/mm^2^) in the SGZ of *Atg7* conditional knockout (cKO; TX) and control (Oil) P21 mice. **g**. Percentage of P13 active RGLs (BrdU^+^) entering quiescence (BrdU^+^MCM2^−^, red) or remaining active (BrdU^+^MCM2^+^, purple) in the SGZ of *Atg7* cKO (TX) and control (Oil) P21 mice. **h**. Quantification of quiescent RGLs that become active (BrdU^−^Ki-67^+^ RGLs/mm^2^) in the SGZ of *Atg7* cKO (TX) and control (Oil) P21 mice. **i**. Quantification of total RGLs (SOX2^+^GFAP^+^ RGLs/mm^2^) at P21. **j**. Schematic representation illustrating protein aggregate dynamics and the regulation of the autophagy-lysosomal pathway in the switch between the active (A) and the quiescent (Q) NSC states. **k**. Schematic representation illustrating the proposed *in vivo* role of autophagy in the acquisition and maintenance of dentate gyrus RGL quiescence during the postnatal period. The autophagy gene *Atg7* is cell-intrinsically required for the proper regulation of quiescence in RGLs when they are being spared as a dormant NSC reservoir. Qrev, quiescent NSCs that revert to the active state. Data are presented as mean ± SEM from n≥3 mice. Statistics: unpaired t-test, two tailed. *p<0.05; **p<0.01; ***p<0.001. Representations shown in a., c., f., h., j. and k. were created with BioRender.com.

To confirm this, we next explored the overall expression profile of ALP-related genes in Q-NSPCs and A-NSPCs by bulk RNAseq. In Q-NSPCs, we found a clear enrichment in genes ascribed to the Macroautophagy and Lysosome Gene Ontology (GO) biological functions (Fig. 3a, Table 1). The gene set partly overlapped with ALP-related genes upregulated in *Hopx*-expressing dentate gyrus precursors when they transition from the proliferative state (P4) to the adult quiescent state^4^ (Fig. 3b, Table 1). It was also shared in part by Q-NSPCs acutely isolated from the adult V-SVZ niche^10^ (Extended Data Fig. 6a). A selection of genes was further validated by RT-qPCR in independent samples (Fig. 3c), including genes involved in the initial induction steps of the autophagy pathway (ULK1-complex genes and the mTOR repressor TSC1 gene, Fig.3c, and AMPK genes, Extended Data Fig. 6b) and key downstream components such *Atg7*, whose product is critically required for generating lipidated LC3-II that in turn participates in autophagosome formation (Fig. 3c). AMPK senses the energy status (AMP:ATP ratio) of the cell and is a major upstream positive regulator of autophagy^21^. AMPK was phosphorylated at Thr172 and was thus activated in Q-NSPCs (Fig. 3d). Phosphorylation of its target ULK1 at Ser555 was increased in Q-NSPCs (Fig. 3e). Moreover, raptor phosphorylation at Ser792 was also enhanced (Extended Data Fig. 6c), suggesting an inhibition of the mTOR pathway (a negative regulator of autophagy), further demonstrating that Q-NSPCs are engaged in autophagy induction. Together, the data suggest that Q-NSPCs accumulate protein aggregates of unknown origin and that this is not due to arrested protein degradation. Rather, Q-NSPCs activate AMPK/ULK1 and engage in a transcriptional programme that enhances the autophagy machinery to deal with the protein aggregates.

**Table 1.**
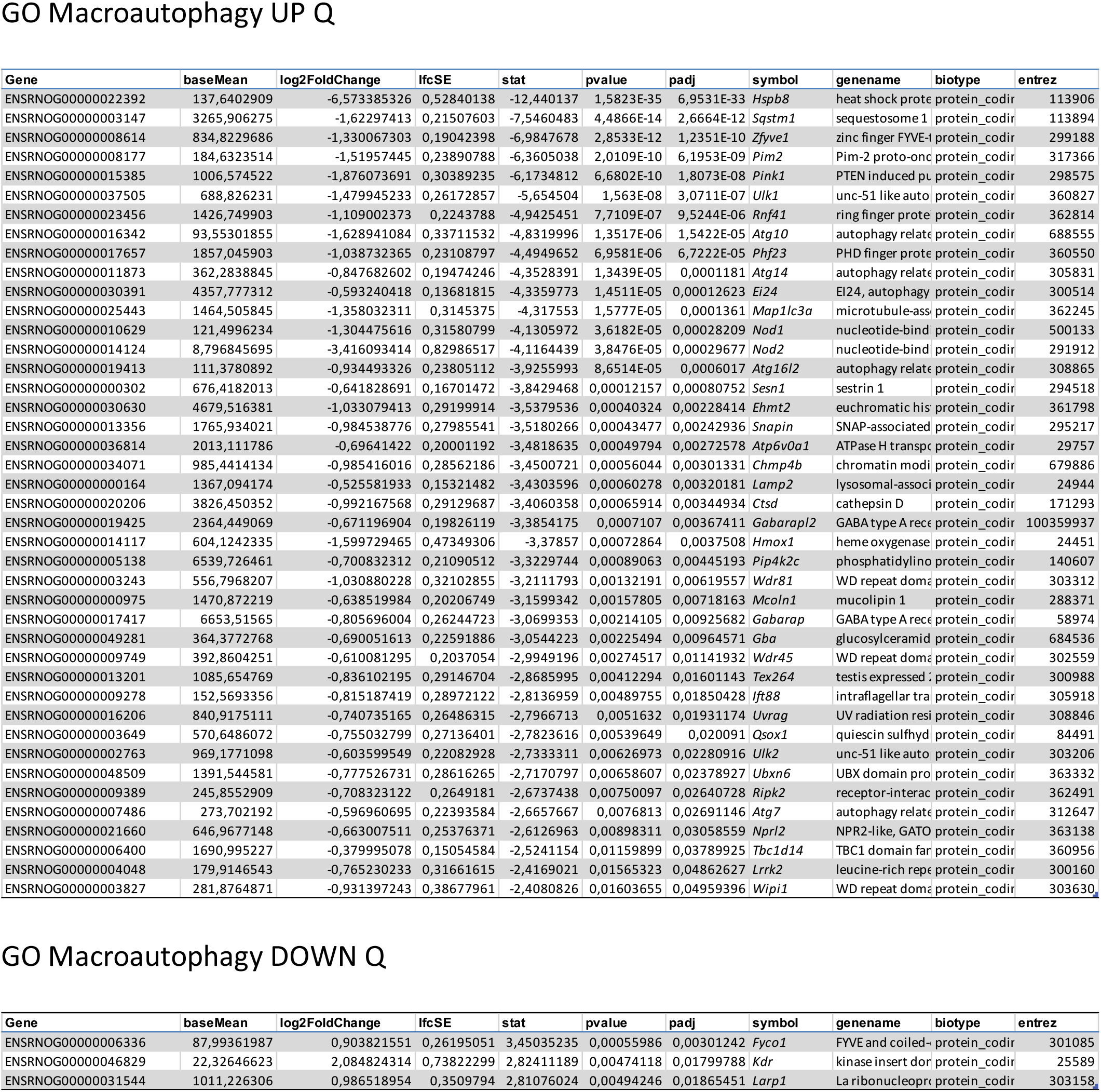
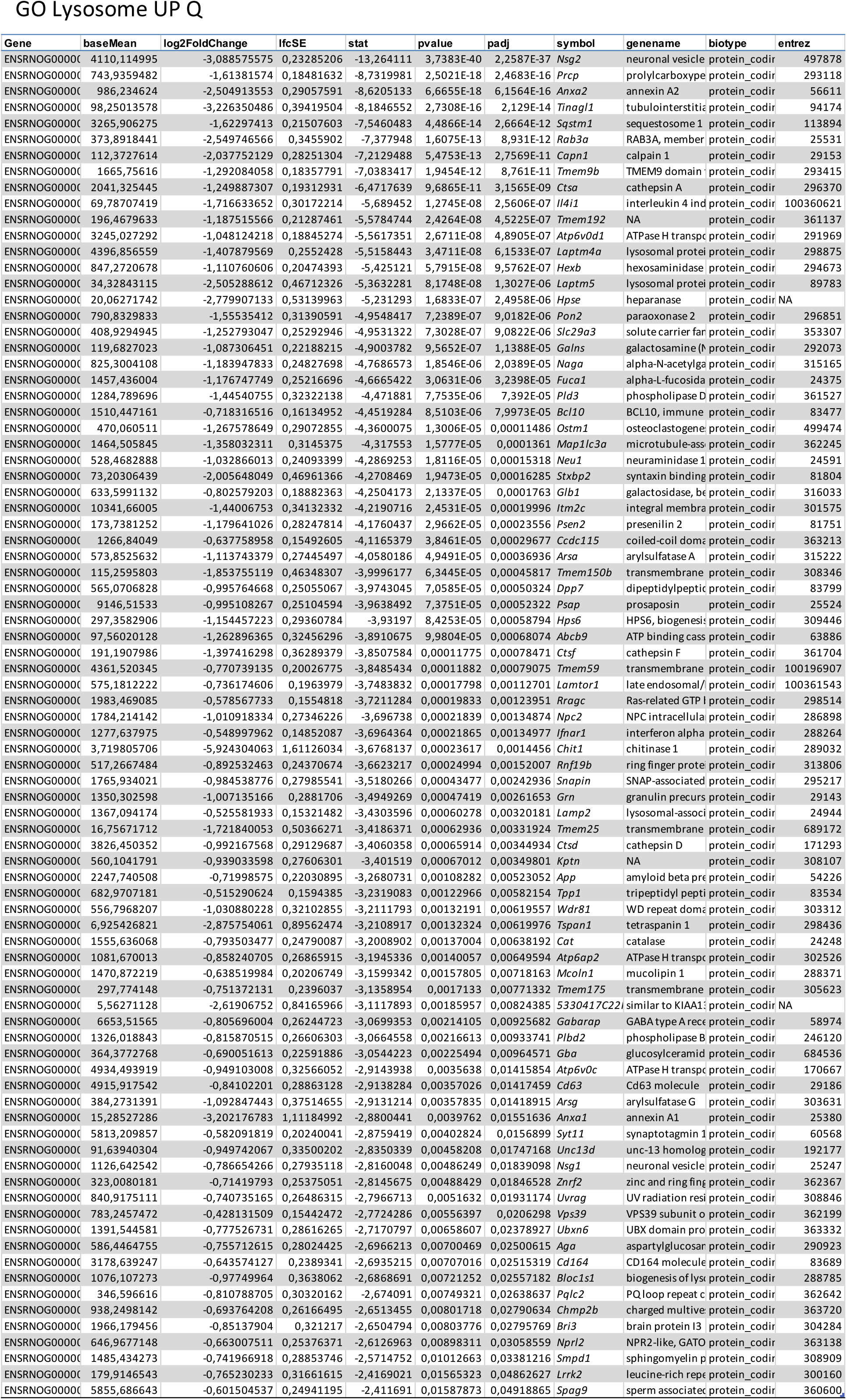
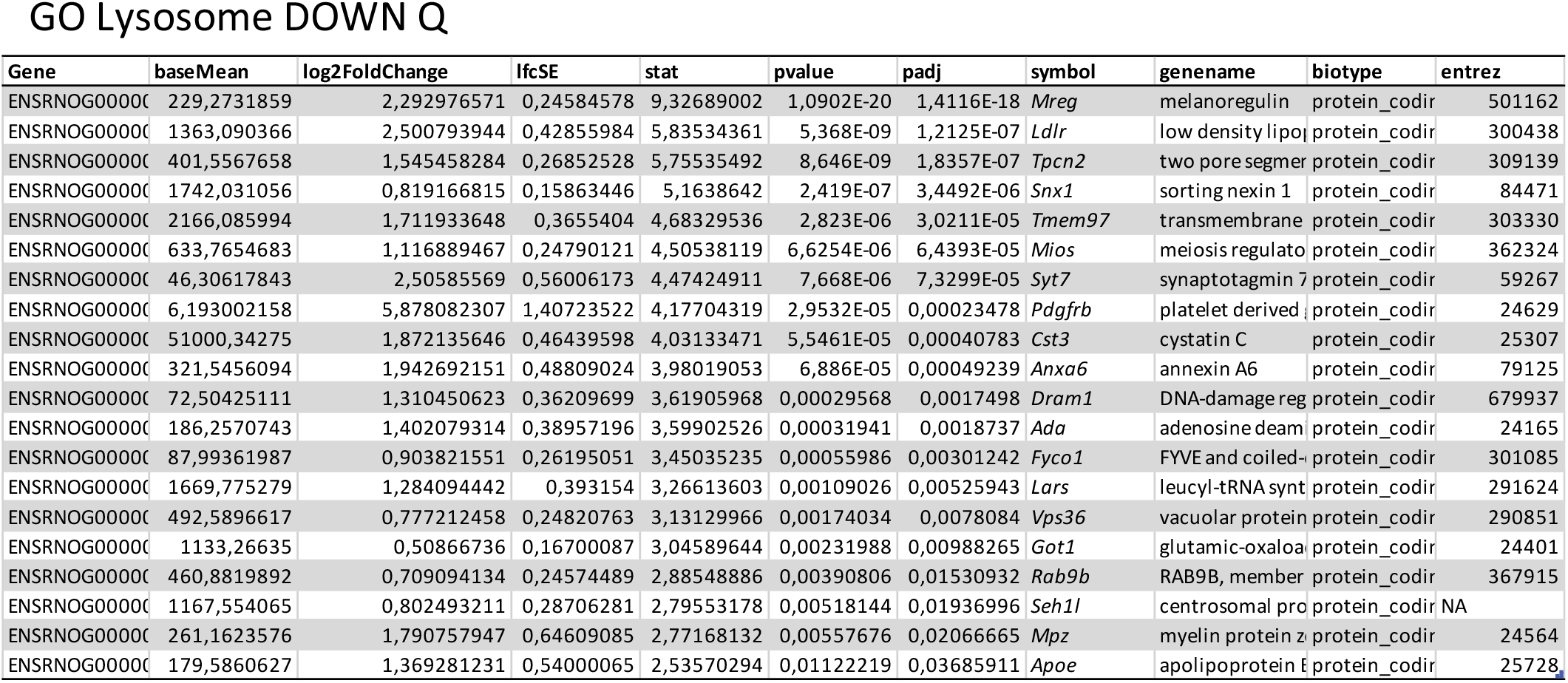
This file contains data for the genes genes ascribed to the Macroautophagy and Lysosome Gene Ontology (GO) biological functions significantly up-regulated (UP) or down-regulated (DOWN) in quiescent NSPCs (Padj<0.05) compared to active NSPCs.

Next we designed a series of *in vitro* experiments to evaluate if autophagy had a positive impact on the entry and maintenance of NSPC quiescence. We employed pharmacological activators and inhibitors of autophagy and identified proliferative cells based on Ki-67 immunostaining (Fig. 4). The AMPK activator phenformin (Phen; Extended Data Fig. 6d) favoured cell cycle exit in NSPC cultures treated with the quiescence inducer BMP4 (Fig. 4a; see Extended Data Fig. 6e for similar proliferation results based on BrdU incorporation). Most importantly, Phen blocked proliferation of A-NSPCs grown in the presence of the mitogen FGF2 and was sufficient to induce cell cycle exit even in the absence of the quiescence-promoting signal (Fig. 4b). Since AMPK has multiple functions beyond regulating autophagy, we employed a selective ULK1 kinase inhibitor (SBI-0206965) to clarify if the Phen effect was dependent on the initiation of the autophagy pathway. Our results demonstrated that instruction of the quiescent state upon increasing AMPK activity was ULK1-dependent (Fig. 4b). Phen also prevented reactivation of the Q-NSPCs when BMP4 was removed (Fig. 4c), whereas the autophagy inhibitor BafA1 favoured cell cycle entry (Fig. 4d).

Finally, we explored the *in vivo* requirement of autophagy in the establishment of the quiescent hippocampal SGZ stem cell reservoir during postnatal development, using a genetic approach. We designed a cell type- and time-specific strategy to accurately examine the role of autophagy in this process. We crossed mice expressing in radial glia–like NSCs a tamoxifen-inducible form of Cre recombinase under nestin transcriptional control^22^ (*NesCreER*^*T2*^), with conditional-knockout mice for the autophagy gene *Atg7*^23^ (*Atg7*^*Fl/Fl*^; Fig. 5a and Extended Fig. 7a-f). To assess the early phase of postnatal SGZ niche development, tamoxifen or oil (control) was administered to P3-P4 animals to ablate *Atg7* in SGZ RGLs. Proliferating cells were labelled by BrdU injection at P5. Entry into quiescence of the actively dividing BrdU^+^ RGLs was evaluated at P14 (a critical timepoint when the morphological pattern and organization of the SGZ niche and DG subfields emerges^3,4^), using the cell cycle marker Ki-67 (Fig. 5a, upper pannel). RGLs were identified throughout the study as SOX2/GFAP co-expressing cells with radial morphology (Fig. 5b). Active RGLs transitioning into quiescence were identified as BrdU^+^Ki-67^-^. We found a robust reduction in the number and percentage of proliferating P5 BrdU^+^ RGLs entering quiescence at P14 in tamoxifen-treated animals (BrdU^+^Ki-67^−^ RGLs, Fig. 5c-d). We also found a significant increase in the total number of P14 RGLs (Fig. 5e), suggesting that proliferating RGLs that fail to acquire dormancy expand through symmetric divisions in the timeframe analysed. As a control, recombination of the targeted *Atg7* allele in the DG after tamoxifen administration was validated by PCR (Extended Data Fig. 7b). Loss of *Atg7* expression in radial glia-like NSCs at the timepoint of evaluation (P14) was corroborated at the mRNA level by RT-PCR of FACS isolated GLAST^+^PROM-1^+^ cells (Extended Fig. 7g).

We also analysed the *Atg7* conditional-knockout mice at a later developmental timepoint, to evaluate the requirement of autophagy in the maintenance of the quiescent stem cell reservoir upon the appearance of the SGZ niche organization. Tamoxifen or oil (control) was administered to P10–P13 animals to ablate *Atg7* in SGZ RGLs and proliferating cells were labelled by BrdU injection at P13. Entry into quiescence of the actively dividing BrdU^+^ RGLs was evaluated at P21 using Ki-67 and MCM2 (Fig. 5a, lower pannel). Similarly, quiescence maintenance was evaluated in BrdU^-^ RGLs (Fig. 5a-b; Extended Data Fig. 8a-b). We found a substantial reduction in the number and percentage of proliferating P13 RGLs that were still entering quiescence at this later postnatal development phase in tamoxifen-treated animals (BrdU^+^Ki-67^−^/BrdU^+^MCM2^−^ RGLs, Fig. 5f-g) and a concomitant increase in the activation of the already quiescent BrdU^−^ RGLs (Fig. 5h). We found no significant changes in the total number of RGLs (Fig. 5i), suggesting that proliferating P21 RGLs, as a population, are not dividing symmetrically and are not being lost through differentiation in the timeframe analysed. As an additional control, we also evaluated *NesCreER*^*T2*^ mice (Extended Data Fig. 8c). We found no differences neither in the entry into quiescence of the actively dividing BrdU^+^ RGLs nor in the activation of the already quiescent BrdU^−^ RGLs when comparing oil *vs*. tamoxifen treated animals (Extended Data Fig. 8d-e). Thus, the results indicate that *Atg7* is cell-intrinsically required for the proper acquisition and maintenance of RGL quiescence in postnatal weeks 1–3, when they are being spared as a dormant NSC reservoir.

## Discussion

Despite extensive literature on the intrinsic and extrinsic cues regulating NSCs in the adult mammalian brain^5^, the mechanisms that cause the initial establishment of NSC quiescence during development remain largely unexplored. Pro-quiescence niche-derived extrinsic signals maintain adult NSCs in a resting state; however, developmental NSCs are set aside as dormant reservoirs before the assembly of a functional niche structure. In addition, most proliferating embryonic and postnatal NSCs do not persist in the mature brain, but those that are being set aside as quiescent, likely need to be equipped with efficient proteostatic mechanisms for proper quality control and turnover of cytoplasmic contents, to ensure their lifelong survival as non-mitotic cells. In this manuscript we demonstrate that autophagy plays a fundamental cell-intrinsic mechanistic role in the developmental transition from NSC proliferation to quiescence.

We focused on the formation of the hippocampal dentate gyrus SGZ NSC niche, which is protracted compared to the development of other brain structures, occurring mainly after birth between postnatal day P3 and P21. We found that dentate gyrus SGZ radial-glia like NSCs experience a proteostatic shift during the postnatal switch from proliferation to quiescence. This shift is evidenced by a reduction in protein synthesis, accumulation of protein aggregates, increased expression of ALP-related genes and higher lysosomal content. It is recapitulated *in vitro* in NSPC cultures challenged with the quiescence-promoting signal BMP4 (Fig. 5i). We further demonstrate that selective ablation of *Atg7*, a key gene for autophagosome formation, in hippocampal radial glia-like NSCs at early and late postnatal stages compromises, respectively, the initial acquisition and the maintenance of quiescence during dentate gyrus morphogenesis (Fig. 5j). Thus, autophagy is critically required by RGLs at the time they are being spared as a dormant pool in the emerging SGZ niche.

Previous studies in adult NSCs focused on protein aggregate accumulation and/or lysosomal function, paying little attention to the autophagy phase of ALP. The accumulation of protein aggregates and lysosomal structures experienced by hippocampal GLAST^+^PROM-1^+^ NSCs and RGLs during development (uncovered in this study) is shared with that reported for adult and aged NSCs from the mammalian V-SVZ neurogenic niche^10^. However, we have encountered discrepancies in the protein homeostasis network employed by the cells that could be attributed to niche or stage differences.

First, adult active V-SVZ NSCs exhibit increased expression of UPS-associated genes and higher proteasome activity^10^; by contrast, in the cell culture hippocampal NSPC model employed in this study we did not detect differences in proteasome activity between the active and quiescent states. We nevertheless found a clear and significant increase in a variety of autophagy markers in quiescent cells. The raise in ProteoStat signal and the accumulation of LC3-II, SQSTM/p62, P-AMPK and P-ULK1 along with the aggrephagy receptor Tax1bp1 and LAMP2 suggests that autophagy increases in Q-NSPCs possibly as a compensatory response to the protein aggregates. Indeed, AMPK/ULK1 activation by phoshorylation clearly indicates that Q-NSPCs are engaged in autophagy induction. At a global scale, in our cell culture conditions, A-NSPCs and Q-NSPCs possibly deal similarly with short-lived soluble proteins that are preferentially degraded through the UPS, while Q-NSPCs likely employ the autophagy pathway to cope with the degradation of long-lived protein aggregates that cannot be diluted through cell division. Uncovering the nature of the protein aggregates or exploring additional roles of the autophagy vesicles in other forms of selective autophagy, such as mitophagy or ERphagy, are beyond the scope of this study, which aimed at elucidating the functional impact of autophagy in the developmental transition from NSC proliferation to quiescence.

Second, previous work showed that adult quiescent V-SVZ NSCs exhibit increased expression of lysosome-associated genes and enlarged lysosomes but less LC3 accumulation in response to inhibitors of lysosomal acidification, pointing to a slower degradation of autophagosomes by lysosomes^10^. Instead, in our system Q-NSPCs had higher content not only in lysosomes, as reported for adult V-SVZ NSCs, but also in many active components of the autophagy machinery, accumulated higher LC3-II levels in response to BafA1 and had a similar autophagic flux compared to A-NSPCs in the timeframe analysed. A similar flow along the vesicular pathway but higher basal content in autophagic machinery may yield higher net autophagy levels per cell in the quiescent sate. Since autophagic flux is an accepted proxy for autophagic degradation activity, our results are in accordance with a recent report that employed NSC cultures from the adult V-SVZ and SGZ^13^, showing LC3 accumulation and higher cathepsin activity (which reflects enhanced lysosomal function) in quiescent cells.

Additional important issues arise when comparing our *in vitro* functional assays with previously published studies performed on adult NSC cultures that reached partially contradictory conclusions. Leeman et al. manipulated ALP with BafA1, which blocks lysosomal acidification, and found decreased activation of the quiescent V-SVZ NSCs in the presence of EGF and FGF2^10^. On the contrary, Kobayashi et al. found increased activation of NSCs with BafA1, which was related to the blockade of activated P-EGFR degradation through the lysosome in an autophagy-independent manner^13^. The decreased proliferative activity upon lysosome accumulation in adult NSCs was linked mechanistically to EGF receptor endocytosis and endolysosomal degradation, not to the role of lysosomes in ALP^13^. In line with Kobayashi et al.^13^, our cell culture results also support higher activation of Q-NSPCs with BafA1 treatment. Nevertheless, all the assays in the current study were performed in the absence of EGF (only FGF2 was used as a mitogen) and thus are likely unrelated to EGFR signalling. Importantly, compared to previous studies, our data uncover the instructive capacity of enhancing autophagy through AMPK/ULK1 signalling for the entrance into quiescence. We demonstrate that increasing AMPK activity with Phenformin is sufficient to promote cell cycle exit of proliferating NSPCs in an ULK1-dependent, and therefore autophagy-dependent, manner.

The RNAseq data presented in this study for cultured NSPCs and hippocampal NSCs during development also demonstrate that stem cells engage in a transcriptional programme that enhances the expression of both autophagy and lysosomal genes when they transition from the proliferating to the quiescent state. Finally, the *in vivo* analysis of *Atg7* conditional knockout mice allowed us to accurately examine the cell-intrinsic role of autophagy in NSCs of the developing hippocampal dentate gyrus at different stages. Altogether our results unfold a novel role for autophagy in the initial establishment and maintenance of radial glia-like NSC quiescence during dentate gyrus SGZ niche formation. The current data support a model wherein autophagy acts as a cell-intrinsic driver in the onset of RGL quiescence during early postnatal development and as a proteostatic pathway involved in the maintenance of the quiescent state once fully acquired at later postnatal stages (Fig. 5j). Both steps are key for the preservation of hippocampal stem cells and neurogenesis throughout adulthood. In summary, our study highlights the involvement of proteostasis and autophagy in the acquisition of NSC dormancy during brain development. We largely extend recent findings in mice on the role of lysosomes in adult quiescent NSC reservoirs^10,13^ and pave the way for further studies aiming to elucidate the mechanistic details of the developmental acquisition of quiescence by the privileged stem cells that populate the mature mammalian brain.

