## Extended Data and Online Methods for "Autophagy is a cell-intrinsic driver of neural stem cell quiescence in hippocampal dentate gyrus development"

### Extended data Figure 1

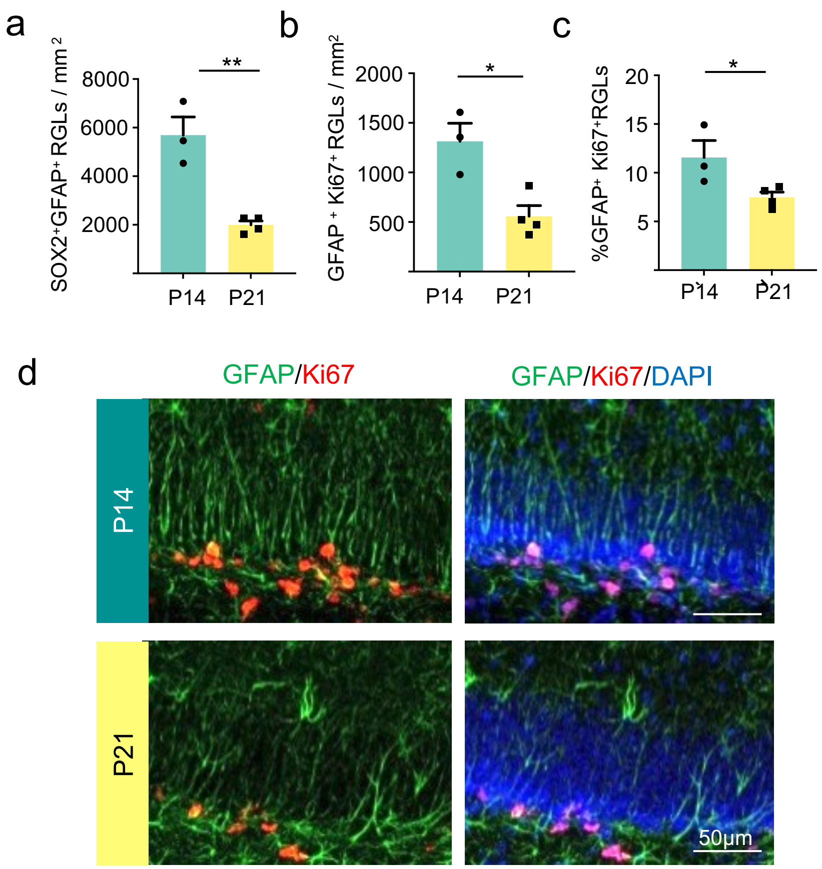


**Extended data Figure 1. Dentate gyrus P14 RGLs continue to enter quiescence until P21. a.** Quantification of RGLs ($GFAP^+/SOX2^+$)/mm2 in the dentate gyrus of postnatal 14 (P14) and 21 (P21) days old wild type mice. **b.** Quantification of active RGLs ($Ki67^+/GFAP^+$)/mm2 in P14 and P21 mice. **c.** Percentage of active RGLs in P14 and P21 mice. Data are represented as mean ± SEM from n≥3 mice. Statistics of a, b, c: Unpaired t-test, two-tailed. **d.** Representative immunohistochemistry confocal images of RGLs labelled for GFAP (green) and Ki67 (red). p-value: *<0.05; **<0.001.

### Extended data Figure 2

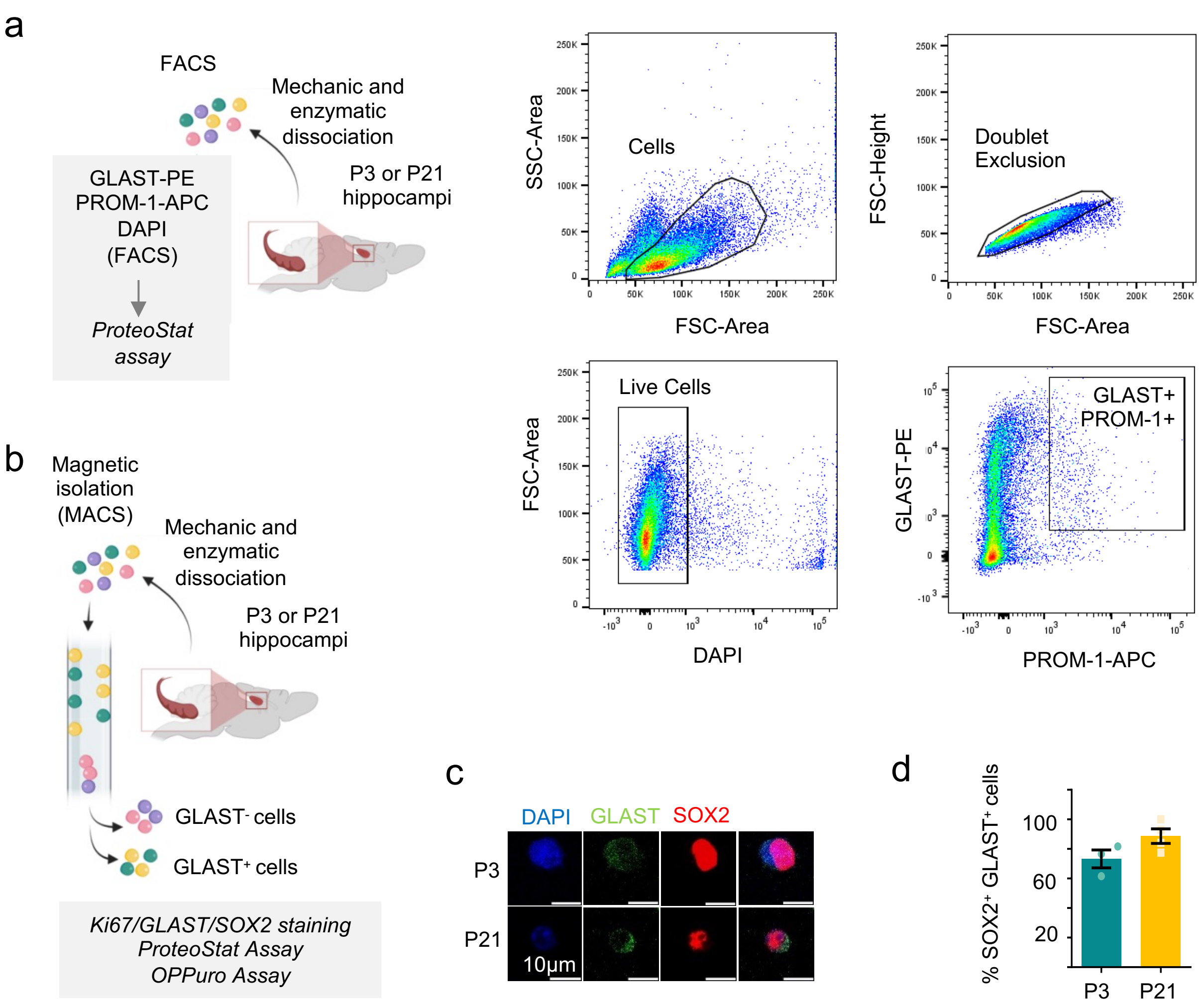


**Extended data Figure 2. NSC isolation and characterization by FACS and MACS. a.** (Left) Schematic representation of NSCs isolation by FACS from postnatal 3 (P3) and 21 (P21) day old mice hippocampi using GLAST-PE and PROM-1-APC antibodies. (Right) Representative images of FACS gating strategy (P3 mice). Forward scatter/side scatter gatings were used to remove doublets and debris. DAPI negative cells were considered viable. NSCs were isolated based on GLAST-PE and PROM-1-APC positive expression. NSCs sorted from P3 and P21 mice were 2.16 ± 0.53 and 0.54 ± 0.13% (mean ± SEM), respectively, of the total cells analyzed. **b.** Schematic representation of magnetic GLAST$^{+}$ cells isolation from postnatal 3 (P3) and 21 (P21) days old mice hippocampi using MACS (Miltenyi Biotec) disaggregation and isolation kit. **c.** Representative immunocytochemestry images of GLAST/SOX2 in GLAST$^{+}$ cells isolated of postnatal 3 and 21-day-old mice. **d.** Percentage of SOX2$^{+}$GLAST$^{+}$ cells isolated from P3 and P21 mice. Data are represented as mean ± SEM from ≥3 independent isolations. Statistics: unpaired t-test, two-tailed. Both representations in a. and b. were created with BioRender.com.

### Extended data Figure 3

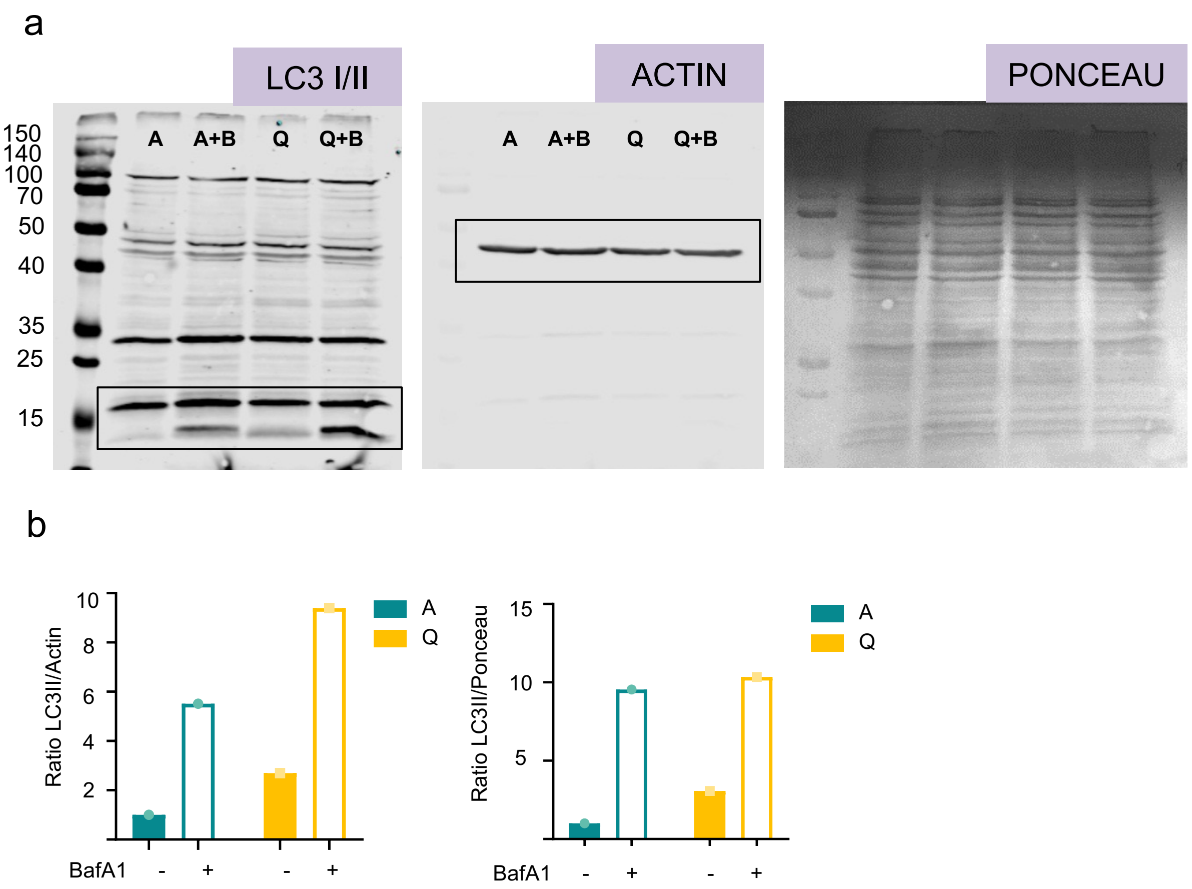


**Extended data Figure 3. Autophagy marker LC3 II accumulates in Q-NSPCs. a.** Original full-length Western immunoblots showing the cropped region used to compose the final image displayed in Figure 2f. **b.** Quantification of LC3 II immunoblot in A-NSPCs and Q-NSPCs treated with BafA1 during 6h. β-Actin (left) and Ponceau (right) were used as a loading control to normalize LC3 II levels, respectively.

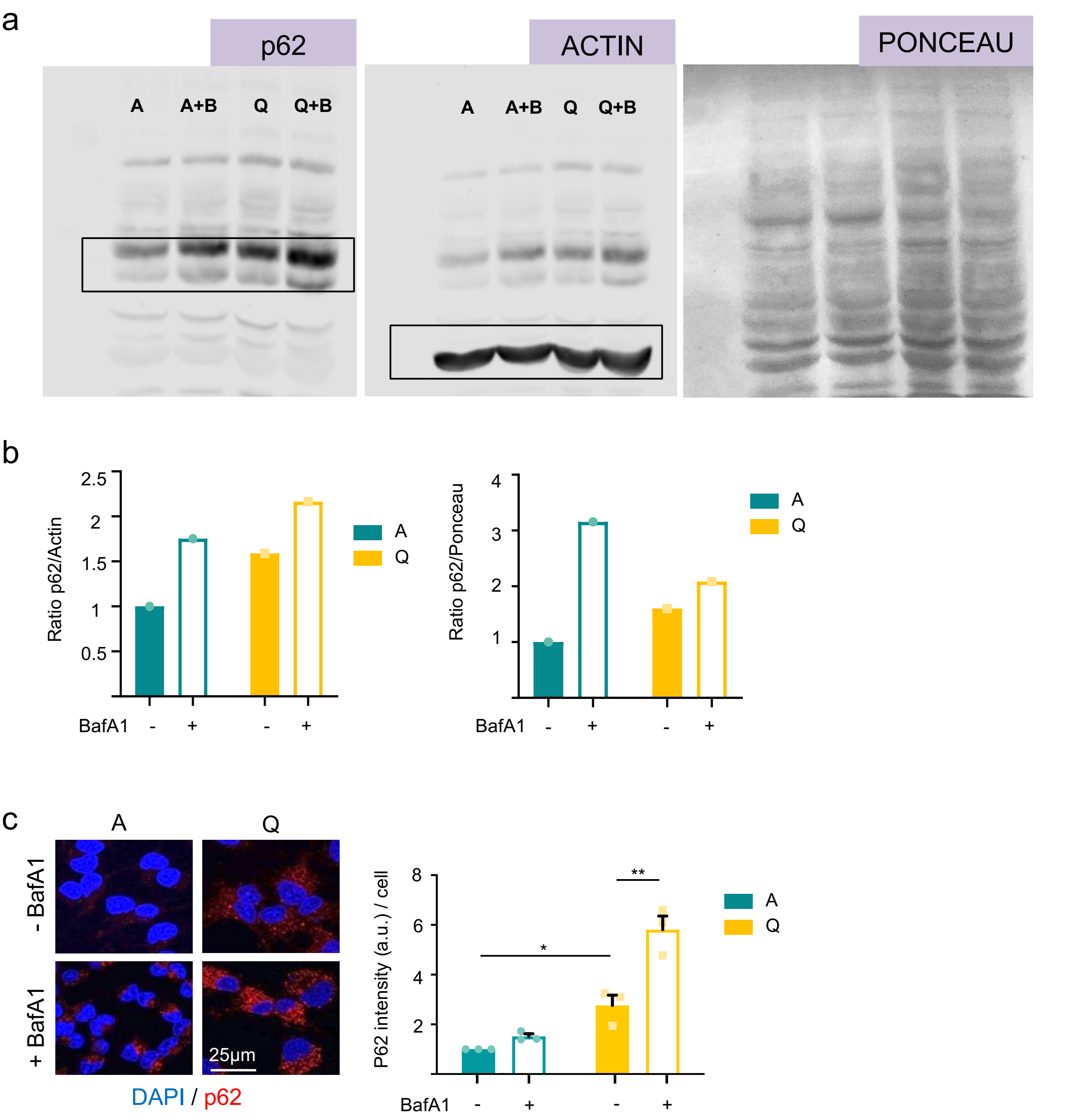


**Extended data Figure 4. Autophagy marker SQSTM/p62 accumulates in Q-NSPCs. a.** Original full-length Western immunoblots showing the cropped region used to compose the final image displayed in Figure 2k. **b.** Quantification of p62 immunoblot in A-NSPCs and Q-NSPCs treated with BafA1 during 6h. β-Actin (left) and Ponceau (right) were used as a loading control to normalize p62 levels, respectively. **c.** (Left) Representative immunofluorescence confocal images of p62 label (red) in Q-NSPC and A-NSPC cultures treated with BafA1. (Right) Quantification of p62 protein level in A-NSPCs and Q-NSPCs treated with BafA1 during 6h. measured as fluorescence intensity. Data are represented as mean ± SEM from n=3 cultures. At least 15 cells were analyzed per experiment and condition. Statistics: two-way ANOVA. p-value: *<0.05; **<0.001.

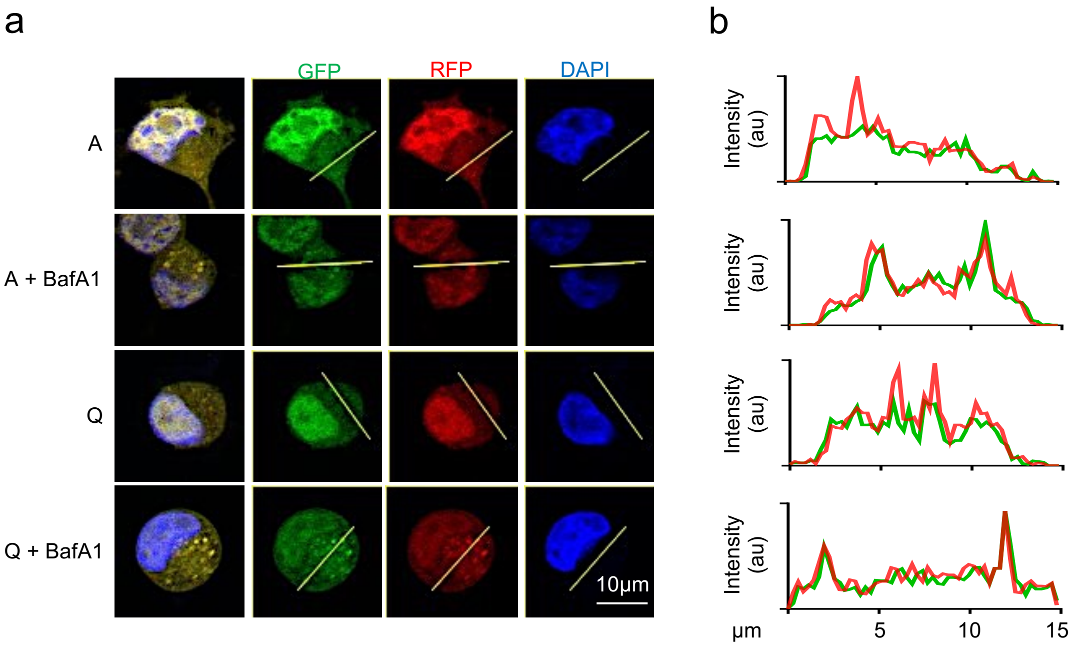


**Extended data Figure 5. NSPC electroporation and pRFP-GFP-LC3 tandem sensor analysis. a.** Representative immunofluorescence confocal images of A-NSPC and Q-NSPC cultures electroporated with pRFP-GFP-LC3 plasmid. Cultures were treated with 100nM Bafilomycin A1 for 6h. **b.** Fluorescence intensity histogram corresponding to the section plotted on the confocal images. Green and red peaks correspond to autophagosomes, and red only peaks correspond to autolysosomes.

### Extended data Figure 6

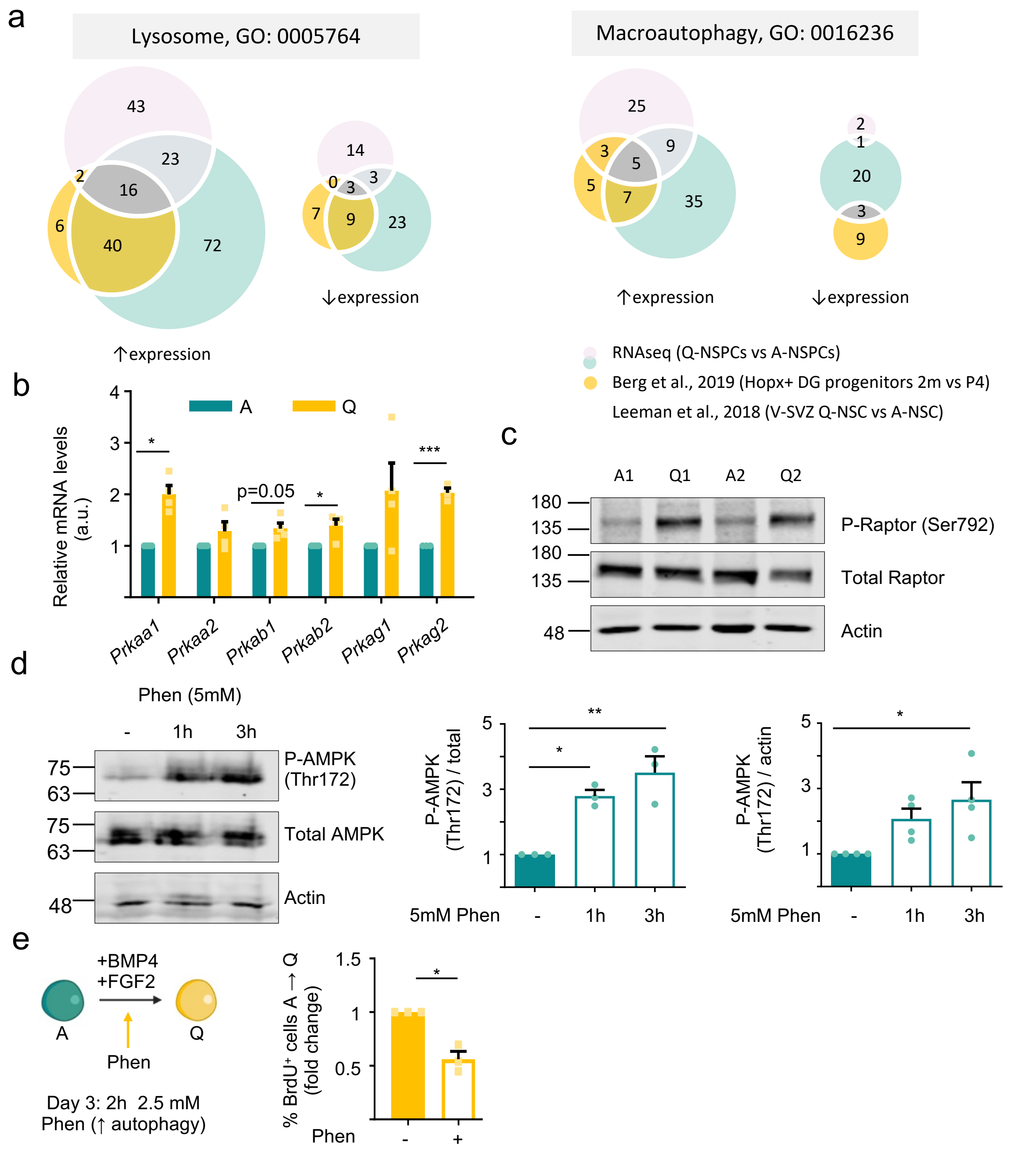


**Extended data Figure 6. Expression of ALP-related genes and Phenphormin assays. a.** Venn diagrams identify commons genes associated to Lysosome GO term (Left) and Macroautophagy GO term (Right) between the RNAseq data from this study, Berg et al., 2019 and Leeman et al., 2018's datasets. **b.** Relative expression of AMPK subunits genes in A-NSPCs and Q-NSPCs. Data are represented as mean ± SEM from ≥3 cultures. Statistics: one-sample t-test. **c.** Representative immunoblots of Raptor and P-Raptor (Ser792) in A-NSPCs and Q-NSPCs. β-Actin was used as a loading control. **d.** (Left) Representative immunoblots of AMPK and P-AMPK (Thr172) in A-NSPCs treated with Phenformin (Phen) during 1 or 3h. β-Actin was used as a loading control. (Right) Quantification of activation of AMPK (P-AMPK Thr172) in A-NSPCs after Phenformin treatment. Data are represented as mean ± SEM from ≥3 cultures. Statistics: 1-way ANOVA. **e.** (Left) Functional assay scheme. Phenformin (Phen) activates the autophagy–lysosome pathway. A-NSPCs were treated for 2 h with Phen on Day 3 of quiescence induction. (Right) Fold-change in the percentage of BrdU$^{+}$ cells 24 h after the 2 h Phen treatment of A-NSPCs entering quiescence. Data are presented as mean ± SEM from n=3 experiments. Statistics: one-sample t-test. p-value: *<0.05; **<0.001; ***<0.0001. Representation shown in e. was created with BioRender.com.

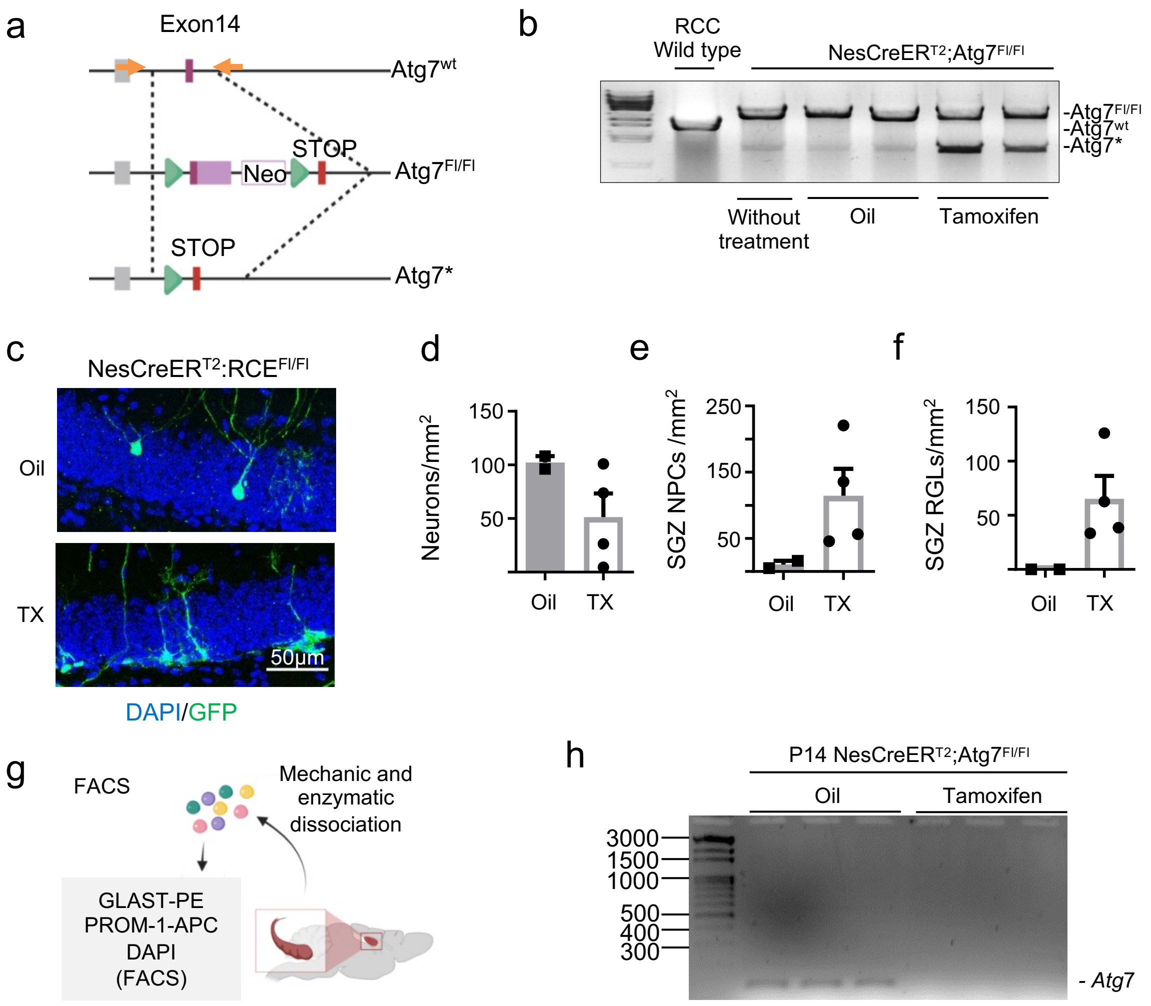


**Extended data Figure 7. Validation of Atg7 allele recombination and loss of Atg7 expression in radial glial-like NSCs of tamoxifen-treated *NesCreER$^{T2}$;Atg7$^{Fl/Fl}$* mice. a.** Schematic representation of the targeted allele of *Atg7* gene. Green triangles indicate loxP sequence. Atg7* represent *Atg7* allele after tamoxifen treatment. Orange arrows indicate position of primers used to corroborate recombination. **b.** Validation of recombination in dentate gyrus after tamoxifen treatment by PCR. **c.** 2 months-old-mice *NesCreERT2;RCE$^{FL}$* were treated with 2mg/day tamoxifen (TX) during 5 days to express GFP in NSCs. Representative immunohistochemistry confocal images of GFP staining in the dentate gyrus of *NesCreERT2;RCE$^{FL}$* 12 days after Tamoxifen treatment. **d, e, f.** Quantification of GFP$^+$, neurons, GFP$^+$ SGZ NPCs and GFP$^+$ RGLs in *NesCreERT2;RCE$^{FL}$*. Data are represented as mean ± SEM from n=2 and n=4 Oil-treated and TX-treated mice, respectively. Statistical analysis was not performed since only n=2 Oil-treated mice were counted. **g.** Schematic representation of NSCs isolation by FACS from postnatal 14 (P14) day old *NesCreER$^{T2}$;Atg7$^{Fl/Fl}$* mice hippocampi using GLAST-PE and PROM-1-APC antibodies. **h.** Agarose gel electrophoresis of RT-qPCR reaction, showing *Atg7* expression in GLAST$^+$PROM-1$^+$ cells isolated from P14 *NesCreER$^{T2}$;Atg7$^{Fl/Fl}$* mice hippocampus treated with oil/tamoxifen. Atg7 ablation was confirmed in tamoxifen treated mice compared to oil animals. Representations shown in a. and g. were created with BioRender.com.

### Extended data Figure 8

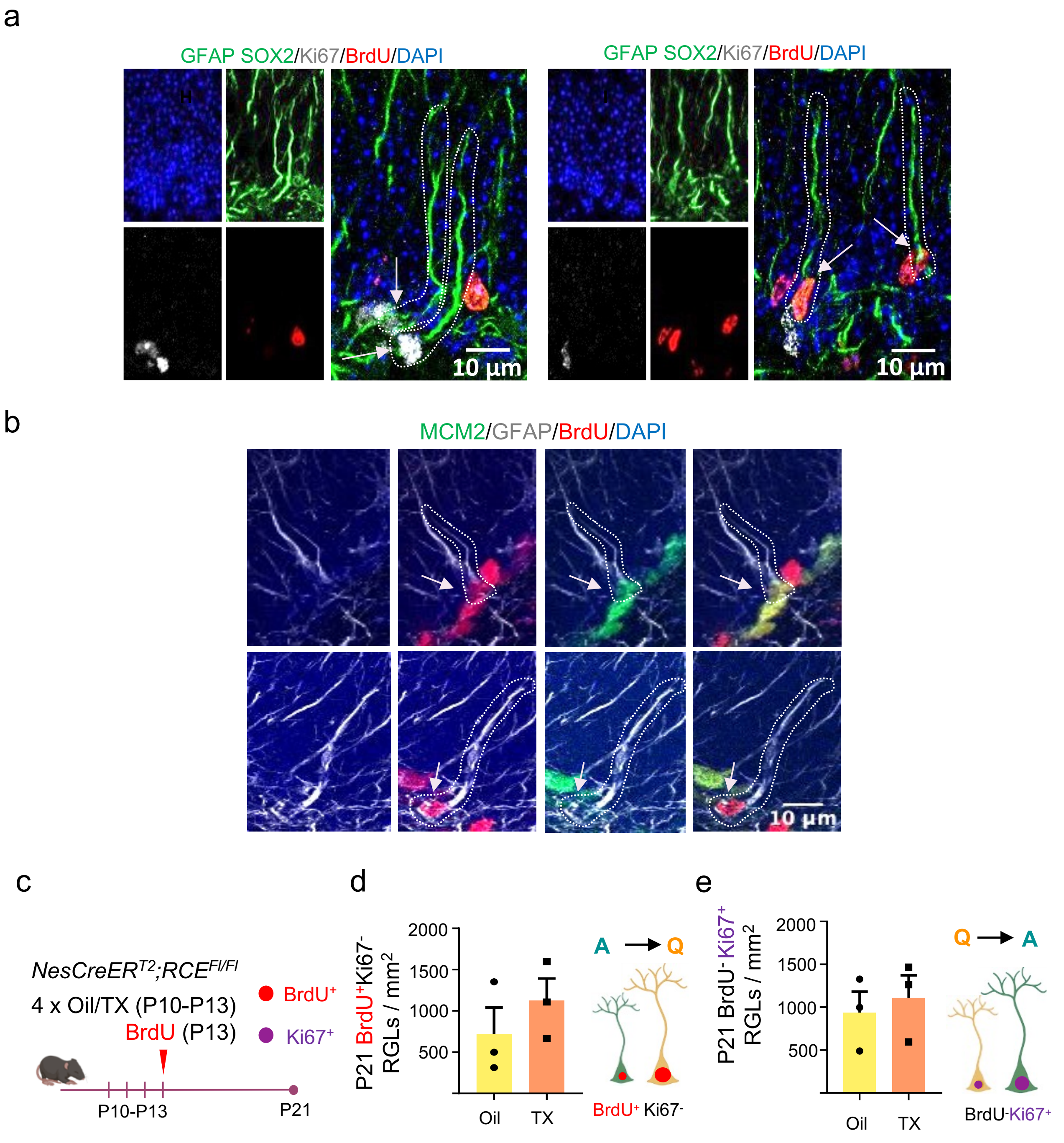


**Extended data Figure 8. RGL analysis in the postnatal SGZ niche and validation of the lack of effect of tamoxifen treatment in *NesCreER$^{T2}$;Atg7$^{Fl/Fl}$* mice. a.** Representative immunohistochemistry confocal images of RGLs labelled for GFAP/SOX2 (green), Ki67 (white) and BrdU (red). $Ki67^{+}BrdU^{-}$ cells (left) and $Ki67^{-}BrdU^{+}$ cells (right) are shown. **b.** Representative immunohistochemistry confocal images of RGLs labelled for MCM2 (green), GFAP (white) and BrdU (red). $MCM2^{+}BrdU^{+}$ cells (above) and $MCM2^{-}BrdU^{+}$ cells (bellow) are shown. **c.** Schematic diagram of the experimental design. A postnatal cohort (P10) of *NesCreERT2;RCE$^{FL/FL}$* mice was injected with 0.25 mg/day tamoxifen (TX) (P10–P13). Oil was used as control. The mice also received 50 mg/kg BrdU injection to mark dividing cells at P13. Animals were sacrificed at P21 and proliferating cells were identified by Ki-67 staining. **d.** Quantification of active RGLs that become quiescent ($BrdU^{+}Ki\text{-}67^{-}$ RGLs/mm$^{2}$) in the SGZ of oil *vs.* tamoxifen treated animals. Data are presented as mean ± SEM from n=3 mice. Statistics: unpaired t-test. **e.** Quantification of quiescent RGLs that become active ($BrdU^{-}Ki\text{-}67^{+}$ RGLs/mm2) in the SGZ of oil *vs.* tamoxifen treated animals. Data are presented as mean ± SEM from n=3 mice. Statistics: unpaired t-test. Representation shown in c. was created with BioRender.com.

**Online Methods**

**Mice**

*NesCreER*$^{T2}$ mice (C57BL/6-Tg(Nes-cre/ERT2)4Imayo) were crossed with *Atg7*$^{Fl/Fl}$ (B6.Cg-ATG7tm1Tchi) or *RCE* mice (Gt(ROSA)26Sortm1.1(CAG-EGFP) to obtain *NesCreER*$^{T2}$*;Atg7*$^{Fl/Fl}$ and *NesCreER*$^{T2}$*;RCE*. For experiments, postnatal (P3 or P10) and adult (2 months-old) mice were used. 0.2 mg tamoxifen was intraperitoneally injected into postnatal mice by one injection per day for two consecutive days (P3-P4) to induce *Atg7* deletion in *NesCreER*$^{T2}$*;Atg7*$^{Fl/Fl}$ mice. 0.25 mg tamoxifen was intraperitoneally injected into postnatal mice by one injection per day for four consecutive days (P10-P13) to induce *Atg7* deletion (*NesCreER*$^{T2}$*;Atg7*$^{Fl/Fl}$ mice) or GFP expression (*NesCreER*$^{T2}$*;RCE* mice). 100 or 50mg/Kg BrdU was intraperitoneally injected into postnatal mice by one injection at P5 or P13, respectively. For control experiments (extended data Fig. 7a-f), 2mg tamoxifen was intraperitoneally injected into 2-month old mice by one injection per day for five consecutive days to induce *Atg7* deletion or GFP expression. Tamoxifen injected mice were housed in individual cages and sacrificed at the times indicated in each experiment. Validation of recombination after tamoxifen treatment was performed by PCR and qPCR (extended data fig. 7b and h, respectively). DNA was extracted from dentate gyrus of 2-month old *NesCreER*$^{T2}$ *Atg7*$^{Fl/Fl}$ mice after tamoxifen/oil treatment. Mice were maintained under specific-pathogen-free conditions and all manipulations were approved by the Animal Care Committee of the Instituto de Biomedicina de Valencia-CSIC.

Animals were anesthetized with pentobarbital (80mg/Kg) and perfused with saline and 4% PFA followed by 2 or 16 h postfixation (postnatal and adult mice, respectively) in 4% PFA at 4°C. Postnatal brains were embedded in 4% agarose solution prior to vibratome processing. The brains were coronally sectioned in a vibratome (Leica Microsystems VT-1200-S). The resulting 40 µm free floating sections were collected sequentially generating antero-posterior reconstructions of the hippocampus conformed by 1 section every 320 µm of hippocampal structure. Stereology was performed by the analysis of at least 3 coronal sections, 40 µm each, separated 320 µm one to each other. Sampling started at first appearance of the infrapyramidal blade of the dentate gyrus.

**MACS and FACS**

GLAST positive cells were isolated by MACS. Hippocampi from postnatal day 3 or day 21 C57Bl6/JRccHsd mice were dissociated using *Neural Tissue Dissociation Kit* (Miltenyi, 130-092-628) with gentle MACS Octo Dissociator (Miltenyi, 130-095-937). Cells were labeled with *Anti-GLAST* (ACSA1) *microBead Kit* (Miltenyi, 130-095-862) and magnetic separation was performed through MS column (Miltenyi, 130-042-102). Cells were attached to 100µg/ml matrigel coated dishes for further analysis. For hippocampal NSCs isolation, cells were labelled with GLAST (ACSA-1)-PE (Miltenyi, 130-118-483) and Anti-Prominin-1-APC (Miltenyi, 130-102-197) antibodies and were isolated by FACS (BD FACSAria™ III Cell Sorter). Compensations were done on single-color controls and gates were set on unstained samples. Dead cells were excluded by staining with DAPI (1 mg/ml). Sorted NSCs were collected directly into N2 medium and plated in 24-well matrigel coated dishes. After 1h, NSCs were fixed with 2% paraformaldehyde and used for subsequent analysis.

For hippocampal NSCs isolation of P14 *NesCreER*$^{T2}$;*Atg7*$^{Fl/Fl}$ mice treated with oil/tamoxifen, cells were labelled with GLAST (ACSA-1)-PE (Miltenyi, 130-118-483) and Anti-Prominin-1-APC (Miltenyi, 130-102-197) antibodies and were isolated by FACS (BD FACSAria™ III Cell Sorter). Sorted NSCs were collected directly into PBS and centrifuged. Pellet was stored at -80ºC until RNA extraction.

**Cell Culture**

For proliferation assays, we used rat Adult Hippocampal Neural Stem and Progenitor Cells (NSPC)[17]. NSPCs were maintained in N2 medium, DMEM/F-12(1:1) (Gibco) adding N2 Supplement (100×) (Gibco), with 10 ng/ml of human fibroblast growth factor 2 (FGF-2) (PeproTech), growing in neurospheres or 20ng/ml of FGF2 growing in poly-ornithine (10 µg/ml)/laminin (5 µg/ml) (Sigma-Aldrich/Millipore) coated dishes[17]. For the quiescence induction, NSPCs were incubated with 50ng/ml BMP4 (PeproTech) and 10ng/ml of FGF2 during 4 days[17]. To revert the quiescence state, NSPCs were incubated again only with N2 medium and FGF2.

To activate autophagy-lysosome pathway NSCPs were treated with 2.5mM of Phenformin (Sigma, 349887) during 2 hours. To specifically inhibit ULK1 before Phenformin treatment, we treated NSPCs with 100 nM of ULK1 kinase inhibitor (Sigma-Aldrich, SBI-0206965) during 1 hour. To inhibit autophagy-lysosome pathway, NSPCs were treated with 50 or 100nM of Bafilomycin (Sigma-Aldrich, B179) during 2 hours. RPF-GFP-LC3 plasmid[20] was transfected into proliferating and quiescent NSPCs by electroporation (10µg plasmid/$10^6$ cells, NEPA21 Super Electroporator). After electroporation, cells were treated with 100nM Bafilomycin during 6h.

**Immunostaining**

Immunohistochemistry and immunocytochemistry were performed using standard procedures. Before fixation with 2% paraformaldehyde (Panreac), cultured neurospheres were disaggregated and seeded in matrigel dishes. Samples (tissue and cells) were incubated with blocking solution (10% Fetal Bovine Serum, 0.2% or 0.5% Triton-X100 (for nuclear or cytoplasmic epitope, respectively) in 0.1M phosphate buffer). Primary antibodies used for the stainings are as follows: Ki67 (Abcam, ab15580), Lamp2 (Biolegend, 108511), p62 (Abcam, ab56416), SOX2 (Gene Tex, GTX101507 and R&D, AF2018), GLAST biotin (Miltenyi, 130-119-161), Tax1bp1 (Novus, NBP3-15794), GFP (Aves-Lab, GFP-1010), GFAP (Sigma, G3893), BrdU (Abcam, ab6326), MCM2 (BD Biosciences, 610700). Treatment with 2N HCl was required before BrdU detection. ProteoStat dye was employed following ProteoStat Aggresome Detection kit instructions (Enzo, ENZ-51035). OPP dye was performed following Click-iT® Plus OPP Protein Synthesis Assay Kits (Molecular Probes, C10456). After staining, cells and all sections were mounted and preserved with 50% Mowiol (Polysciences, 17951), 2.5% DABCO (Sigma, D2522). Images were acquired with a Leica Spectral SP8 confocal microscope with a 40x Oil objective. Images were analyzed with Fiji Image J Software.

**Western blot**

Cell extracts were fractionated by SDS-PAGE and transferred to a polyvinylidene difluoride (PVDF) membrane following the manufacturer's protocol (Bio-Rad). Membranes were incubated with 5% BSA (Sigma) in TBST for 60 min. Primary and secondary antibodies used for the Western blot assays are as follows: LC3 (Santa Cruz, sc-376404), p62 (Abcam, ab56416), P-AMPK (Cell Signaling, 2535), total AMPK (Cell Signaling, 2793), P-ULK (Cell Signaling, 5869), total ULK (Cell Signaling, 8054), P-Raptor (Cell Signaling, 2083), total-Raptor (Cell Signaling, 2280), Actin (Sigma-Aldrich, A5441), IRDye 680LT anti-mouse (Licor, 925-68020), IRDye 800CW anti-rabbit (Licor, 925-32211). Proteins were detected with LI-COR Odyssey and analized with Image Studio™ Lite.

**Gene Expression analysis**

RNA was extracted from cells. cDNA was obtained by reverse-transcription (RT) employing PrimeScrip RT Reagent kit (Takara). Gene expression was determined by quantitative polymerase chain reaction (qPCR) in a QuantStudio 5 (Applied Biosystems) using SYBR PremixEX Taq (2×) (Takara) and the corresponding forward and reverse primer for each gene. *Sdha* was used as the internal reference for normalization. Data were analysed according to the 2−ΔΔCt method. The following primers were employed in qPCR and PCR:

| Gene | Sequence (5'->3') |
|---|---|
| *Atg7* | FW: CTTGACCTTCGCGGACCTAAAGAA |
| | RV: AGGGCCTGGATCTGTTTTGGTGA |
| *Atg7flox* | FW: TGGCTGCTACTTCTGCAATGATGT |
| | RV: TGGACCAAGCCTGGACGTGGACTTT |
| *Hdac6* | FW: AGCCCCAATCTAGCGGAGGTAAAG |
| | FW: AGG GAA GCT GTC ATC CCA AAG GCA |
| *Idua* | FW: TGCCTCACGACCAGGCTGACCA |
| | RV: CGTGAAGTACCCAGAAGGACTGCC |
| *Lamp1* | FW: CCA CAA CTG ACA TCA AGG CAG ACA TC |
| | RV: GTGGGCACAAGTGGTGGTGAGG |
| *Lamp2* | FW: CTG TCT GCT GGC TAC CAT GGG |
| | RV: TGACAGCTGCCGGTGAAGTTGG |
| *Naga* | FW: TCCACGGACCTGCGTACCATCTC |
| | RV: GCATGTCTGTCCTGCGGCTGAAGA |
| *Sdha* | FW: AGAGGACAACTGGAGATGGCATT |
| | RV: AACTTGAGGCTCTGTCCACCAA |
| *Sqstm1* | FW: TTTCAGGCGCACTACCGCGATGA |
| | RV: CGCCGGCACTCCTTCTTCTCTTT |
| *TSC1* | FW: CATGGTGCCTGAAGAAACCA |
| | RV: TGAATTCGCACATGCTCCAT |
| *Ulk1* | FW: CCTTGCCAAGTCCCAGACACTGC |
| | RV: CATAGTGTGCAGGTAGTCAGCCAG |
| *Prkaa1* | FW: ACGTGGTCGGGAAAATCCGC |
| | RV: ACGTCGACTCTCCTTTTCGTCC |
| *Prkaa2* | FW: AGTGAATGCATACCATCTTCGAGTAA |
| | RV: ACCAGACCTCTGCTCCACC |
| *Prkab1* | FW: GGGCACCAAGGATGGGGACAG |

| | |
|---|---|
| | RV: GCTTTCTCATTCACCTCGAGGTCGT |
| *Prkab2* | FW: CTGGCAGCAGGATTTGGATGATTC |
| | RV: ATTATGACTCTTAATCAGAGGGATCTTGG |
| *Prkag1* | FW: ACTTCGCTGCAGGTAAAGAAAGCCT |
| | RV: ATAGATCTGCACCAGGGCTGACTT |
| *Prkag2* | FW: AAGATTGAAACGTGGAGGGAACTGTACT |
| | RV: CATGTCAGACATAAAAAGCTGGAGGAAC |

**Gel electrophoresis**

PCR or qPCR products were resolved in a 2% agarose gel electrophoresis and were visualized under a UV transilluminator.

**RNAseq**

For the transcriptomic analysis employing, total RNA was extracted from active and quiescent NSPCs. cDNA libraries were prepared using the TruSeq Stranded mRNA Library Preparation Kit (Illumina, San Diego, CA). Libraries were sequenced on the Illumina® NSQ 500 High Output KT v2 (75 CYS) (Illumina, San Diego, CA).

**Measurement of proteasomal activity**

*Proteasome 20S Assay kit* (Sigma-Aldrich) was employed to measure chymotrypsin-like activity associated with the proteasome. Active and quiescent NSPCs were grown on poly-ornithine/laminin dishes and 10µM of MG132 was used as control. Proteasome activity was normalized to cell number. For cell number calculations, cells were fixed and stained with crystal violet. $A_{590}$ values were next used to determine cell number by interpolating from a standard curve.

**Statistics**

All statistical tests and sample sizes are included in the Figure Legends. Measurements were taken from distinct samples. Normality was measured with Shapiro Wilks test. Normal data are shown as mean ± SEM and non-normal data are represented as median ± interquartile range. In all cases, p-values are represented as follows: ****p<0.0001, *** p<0.001, ** p<0.01, * p<0.05. For normal data, all quantifications were statistically analyzed using either one sample t-test (normalized data 2 groups), Student's t-test-2 tails (2 groups) or 2 way-ANOVA test. For non-normal data, Wilcoxon Signed Rank Test or Mann Whitney test was employed. Statistical analysis was performed using Graphpad Prism 8. No statistical methods were used to pre-determine sample sizes. Subjects were randomly assigned to treatment groups (Figure 3).

**Data availability**

RNAseq data were registered at the NCBI BioProject and BioSample databases (Bioproject PRJNA666074).

**Acknowledgements**

We thank all members of the Mira laboratory and Prof. Dr. Chichung Lie for discussions and helpful suggestions; J. Durban, R. Viana, M. Quesada and M. Heredia for support with RNAseq, confocal microscopy, histology and animal care, respectively. This work was supported by the Spanish MICINN (grant no. PID2019-111225RB-I00) and PROMETEO/2018/055 grant from Generalitat Valenciana for H.M.; FPI fellowship BES-2016-077156 to I.C-B; and APOSTD2019/058 from Generalitat Valenciana to L.C-C.

**Author contributions**

H.M. conceived the study. I.C-B and H.M. designed the experiments. I.C-B and L.C-C performed the assays, analyzed the data and prepared the figures. I.C-B, J.G-N and C.F-I conducted the postnatal confocal microscopy imaging experiments and analyzed the imaging data. P.S. was involved in the design and interpretation of the autophagy flux and cell signalling experiments. H.M. wrote the main manuscript text. All authors reviewed the manuscript.

**Competing interests**

The authors declare no competing interests.
